# Codon usage and amino acid identity are major determinants of mRNA stability in humans

**DOI:** 10.1101/488676

**Authors:** Megan E. Forrest, Ashrut Narula, Thomas J. Sweet, Daniel Arango, Gavin Hanson, James Ellis, Shalini Oberdoerffer, Jeff Coller, Olivia S. Rissland

## Abstract

mRNA degradation is a critical, yet poorly understood, aspect of gene expression. Previous studies demonstrate that codon content acts as a major determinant of mRNA stability in model organisms. In humans, the importance of open reading frame (ORF)-mediated regulation remains unclear. Here, we globally analyzed mRNA stability for both endogenous and human ORFeome collection mRNAs in human cells. Consistent with previous studies, we observed that synonymous codon usage impacts human mRNA decay. Unexpectedly, amino acid identity also acts as a driver of translation-dependent decay, meaning that primary protein sequence dictates overall mRNA levels and, consequently, protein abundance. Both codon usage and amino acid identity affect translational elongation rate to varying degrees in distinct organisms, with the net result being sensed by mRNA degradation machinery. In humans, interplay between ORF- and UTR-mediated control of mRNA stability may be critical to offset this fundamental relationship between protein sequence and mRNA abundance.

## INTRODUCTION

Messenger RNA (mRNA) decay is critical for post-transcriptional gene regulation. The predominant mechanism of normal cytoplasmic mRNA turnover begins by removal of the 3’ poly(A) tail by the PAN2-PAN3 and CCR4-NOT/CAF1 deadenylase complexes (Wolf and Passmore, 2014; Yamashita et al., 2005). Deadenylation then triggers removal of the 5’ 7-methyl-guanosine cap by the DCP1/2 decapping complex, exposing a free 5’-monophosphate group to the 5’→3’ exonuclease XRN1 and leading to rapid destruction of the transcript body (Beelman et al., 1996; Muhlrad et al., 1994; Yamashita et al., 2005). This deadenylation-dependent decay process is highly conserved from *Saccharomyces cerevisiae* to higher eukaryotes, including mammals.

Mammalian mRNA half-lives vary extensively *in vivo* (Lugowski et al., 2018; Schwanhäusser et al., 2011), and a major question has been which processes and regulatory elements underlie this variation. In mammals, the 3’ untranslated region (UTR) is a hotspot for regulation, containing the majority of instability-promoting sites, such as AU-rich elements or microRNA-binding sites (Bartel, 2009; Rissland, 2016; Shyu et al., 1989; Xu et al., 1998). Typically, instability-promoting motifs accelerate decay by eventually recruiting deadenylation-dependent decay machinery and stripping stabilizing mRNP components (Eulalio et al., 2009; Fabian et al., 2011; 2013; 2009; Kuzuoğlu-Öztürk et al., 2016; Rissland et al., 2017; Zekri et al., 2013). Despite advances, 3’UTR-mediated regulation has so far failed to completely explain the range of half-lives observed in mammalian cells, leaving open the question of whether other parts of the mRNA, such as the open reading frame (ORF) sequence, might modulate mRNA decay rates.

We have previously demonstrated that codon usage is a key mechanism underlying mRNA half-life variability in *S. cerevisiae* such that changes in overall “optimal” and “non-optimal” synonymous codon composition manifest as dramatic changes in mRNA stability (Presnyak et al., 2015; Radhakrishnan et al., 2016). In yeast, optimality is determined by differences in ribosome elongation speed: different synonymous codons are decoded at different rates, primarily due to the balance between tRNA abundance and demand by cognate codons (Gardin et al., 2014; Hanson et al., 2018; Ingolia et al., 2009; Pechmann and Frydman, 2013; Yu et al., 2015). Thus, changing the codon content of a transcript influences not just how quickly elongation proceeds but also its stability with a less optimal transcript being degraded more quickly (Hanson et al., 2018; Presnyak et al., 2015; Radhakrishnan and Green, 2016). Following our initial observations in budding yeast, similar relationships between codon content and mRNA half-life have since been demonstrated in other organisms, including *E.coli* (Boël et al., 2016), *Schizosaccharomyes pombe* (Harigaya and Parker, 2016), *Trypanosoma brucei* (de Freitas Nascimento et al., 2018; Jeacock et al., 2018), *Drosophila melanogaster* (Burow et al., 2018), and zebrafish (Bazzini et al., 2016; Mishima and Tomari, 2016), hinting that this process is broadly conserved. However, the effects of codon optimality and other facets of ORF composition on mRNA have been incompletely explored in higher metazoans.

In this study, we demonstrate that, in humans, both synonymous codon usage and amino acid composition drive a wide range of mRNA half-lives. First, consistent with previous studies (Mattijssen et al., 2017), we found via reporter assays that codon optimality affects mRNA stability. To extend these results transcriptome-wide, we determined half-lives for two sets of transcripts, endogenous mRNAs and those derived from the human ORFeome collection. Because the ORFeome collection expresses human ORFs cloned with invariant UTR sequences, these mRNAs strip away confounding concerns of 3’UTR regulation, differential translational initiation, and other gene-specific effects. As opposed to our previous findings, we found that tRNA abundance failed to explain differences between codons, and, instead, codons encoding the same amino acid seemed to have similar effect on mRNA stability. Subsequently, we identified specific amino acids that can drive mRNA stability; importantly, codons associated with instability-causing amino acids are also translated more slowly. Finally, we found that effects of amino acid composition on mRNA half-life are broadly observed across multiple metazoan species, but not in fungi. Taken together, these findings establish that a combination of codon and amino acid composition drive mRNA stability in metazoans by changing elongation speed, solidifying the importance of the coding sequence itself in controlling gene expression.

## RESULTS

### Codon use affects mRNA stability in human cells

We first sought to assess the relationship between codon optimality on mRNA stability in human cells. We had previously employed species-specific tRNA adaptation index (sTAI) values as a guide for mRNA sequence optimization in *S. cerevisiae*, which primarily relies on tRNA gene copy number as a proxy for tRNA abundance (Radhakrishnan et al., 2016; Sabi and Tuller, 2014). However, human tRNA expression can vary widely in different cell types (Dittmar et al., 2006; Goodarzi et al., 2016; Shigematsu et al., 2017), and so this method can only approximate tRNA abundance in humans. For more accurate optimization, we calculated codon-specific tRNA adaptation index values using published HEK293T tRNA sequencing data (Zheng et al., 2015). “Optimal” and “non-optimal” codons were defined as codons with a tAI greater and less than the median (0.155), respectively.

We designed 11 synthetic firefly luciferase reporters with variable ORF sequences ranging from 0-100% optimal codon content (Figure 1A). Importantly, these constructs differ only in synonymous codon usage, contain the same 5’ and 3’UTRs, and produce identical protein. Optimal codon content correlated with steady-state mRNA abundance such that the 100% reporter was 3.7-times more abundant than the 0% reporter in HEK293 cells (*r_s_* = 0.70; *p* = 0.03, two-tailed t-test; Figure 1B). To determine whether these effects were due to differences in mRNA stability, we performed transcription shutoff via a tetracycline-inducible repressor (Tet-off) system, followed by northern blotting (Figure 1C). Consistent with our steady-state analysis, the 100% reporter was significantly more stable than the 0% reporter (*p* = 0.002, two-tailed t-test; Figure 1C, Figure S1A).

**Figure 1.**
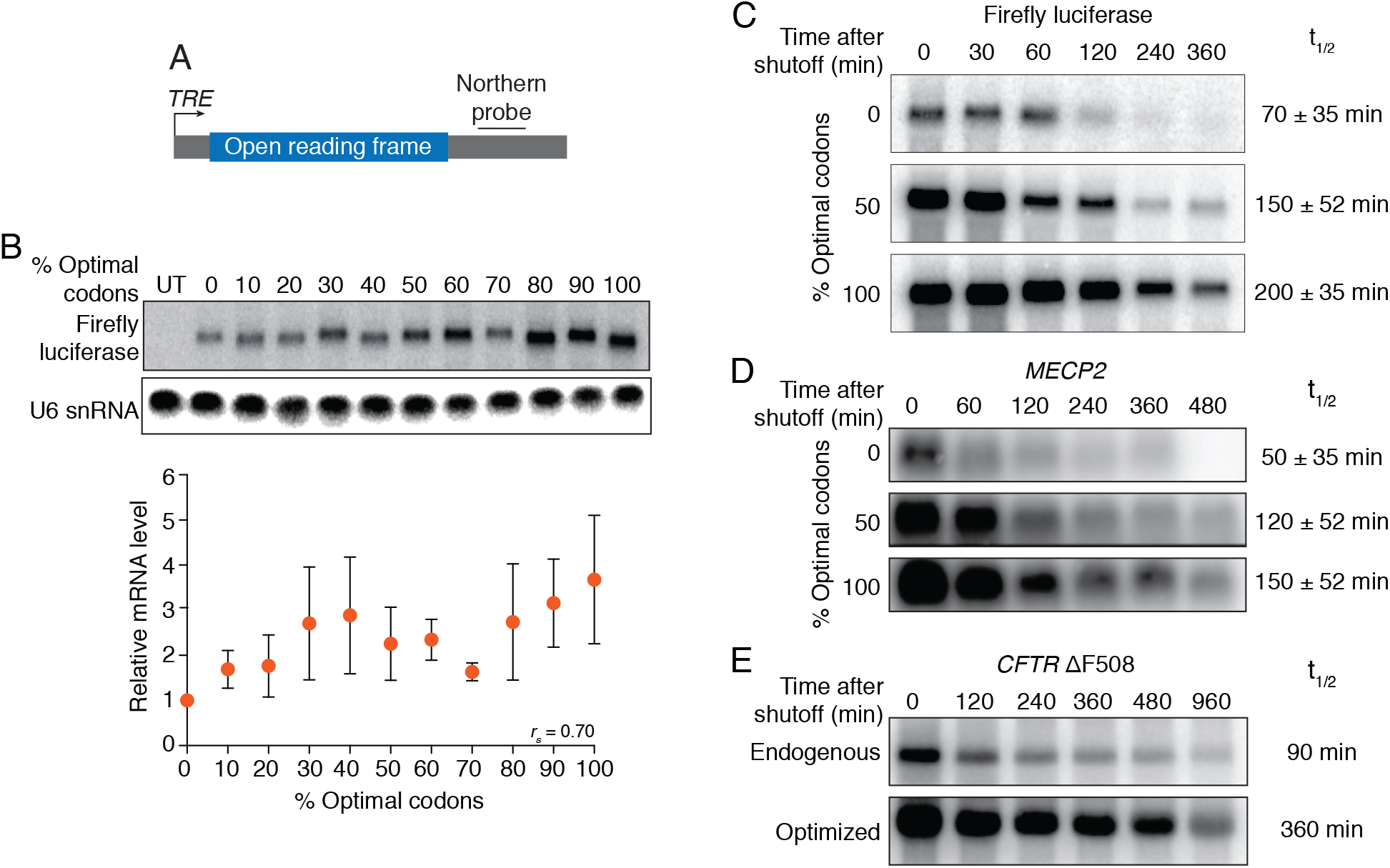
Optimal codon usage modulates mRNA stability in human cells. (A) Steady-state mRNA levels of firefly luciferase variants. Shown is a northern blot, probing for the common 3’UTR within different firefly luciferase reporter mRNAs, which vary by optimal codon content, and U6 snRNA loading control. Quantification of Firefly luciferase mRNA (relative to 0% optimal construct) is shown at bottom (error bars denote standard deviation; n=3). (B) More optimal luciferase mRNA variants are more stable. Transcriptional shut-off experiments were performed in Tet-Off HEK293 cells, and firefly luciferase mRNA levels were determined by northern blots. Timepoints correspond to the time after the addition of doxycycline, which shut-offs transcription of the reporter. t½ corresponds to the half-life (min) ± standard deviation (n=3). See Figure S1 for loading control. (C) More optimal *MECP2* mRNA variants are more stable. As in B, expect for *MECP2*. See Figure S1 for loading control. (D) More optimal *CFTR* ΔF508 mRNA variants are more stable. As in B, except for *CFTR* ΔF508. See Figure S1 for loading control. See also Figure S1 and Tables S1 and S2.

Having observed that codon content affected the stability of a reporter mRNA, we next investigated whether the same was true for endogenous human genes. We looked at effects of altering codon optimality on a human mRNA *MECP2*, which has a relatively high optimal codon content as calculated by tAI (68% optimal codons vs. 59% overall median). Loss-of-function mutations in *MECP2* are the most common cause of Rett syndrome, which is characterized by severe neurological deficits (Liyanage and Rastegar, 2014; Martínez de Paz and Ausió, 2017). As before, we produced reporters of variable optimal codon content and measured their stability. Similar to the firefly luciferase reporters and previous reports on *LARP4* mRNA (Mattijssen et al., 2017), changing *MECP2* codon content altered mRNA stability (*p* = 0.01, two-tailed t-test; Figure 1D, Figure S1B). Finally, we investigated ΔF508 *CFTR* mRNA, which is the most common causative allele for cystic fibrosis (Ferec and Cutting, 2012). The endogenous CFTR ΔF508 mRNA contains an average level of optimal codons as calculated by tAI (62% optimal codons vs. 59% overall median). Optimization of the endogenous sequence (“optimized” CFTR: 85% optimal codons by tAI) resulted in an increase mRNA stability (Figure 1E, Figure S1C). Taken together, these data demonstrate that altering codon optimality (as defined by tRNA abundance) can impact mRNA stability in human cell lines.

### Human mRNA stability is broadly influenced by coding region determinants

Having found that changing codon content altered mRNA stability for a handful of reporters, we next asked whether these trends held on a transcriptome-wide scale. However, in addition to varying in their coding sequences, mRNAs also vary in their 5’ and 3’UTRs. Because UTR-mediated regulation has evolved in conjunction with ORF-mediated regulation, we reasoned that it would be challenging to separate out correlative and causative effects of the coding sequence on mRNA stability. To address this issue, we turned to the human ORFeome collection, which contains ~16,000 full-length ORFs (corresponding to ~14,000 genes) in a lentiviral expression system (Yang et al., 2011). Importantly, the ORFs are flanked by invariant 5’ and 3’ UTRs, as well as a V5 tag, thus allowing us to isolate effects of coding sequences on transcript stability.

We divided the ORFeome collection into six pools and made stable HEK293T cell lines from two (Figure 2A). As determined by western blot, ORFeome-derived proteins were expressed, and pool complexity was maintained through cell line creation (Figure S2A). We next measured half-lives of ORFeome-derived mRNAs using metabolic labeling and approach-to-equilibrium kinetics (Lugowski et al., 2018). Because we prepared RNA sequencing libraries without any enrichment for ORFeome-derived transcripts, we used stringent cut-offs to classify transcripts as ‘endogenous’ or ‘ORFeome-derived’ (see Supplemental Information, Figure S2B, C). For transcripts expressed both endogenously and from the ORFeome, we could not determine the origin of the associated reads, and so these transcripts were excluded in downstream analysis.

**Figure 2.**
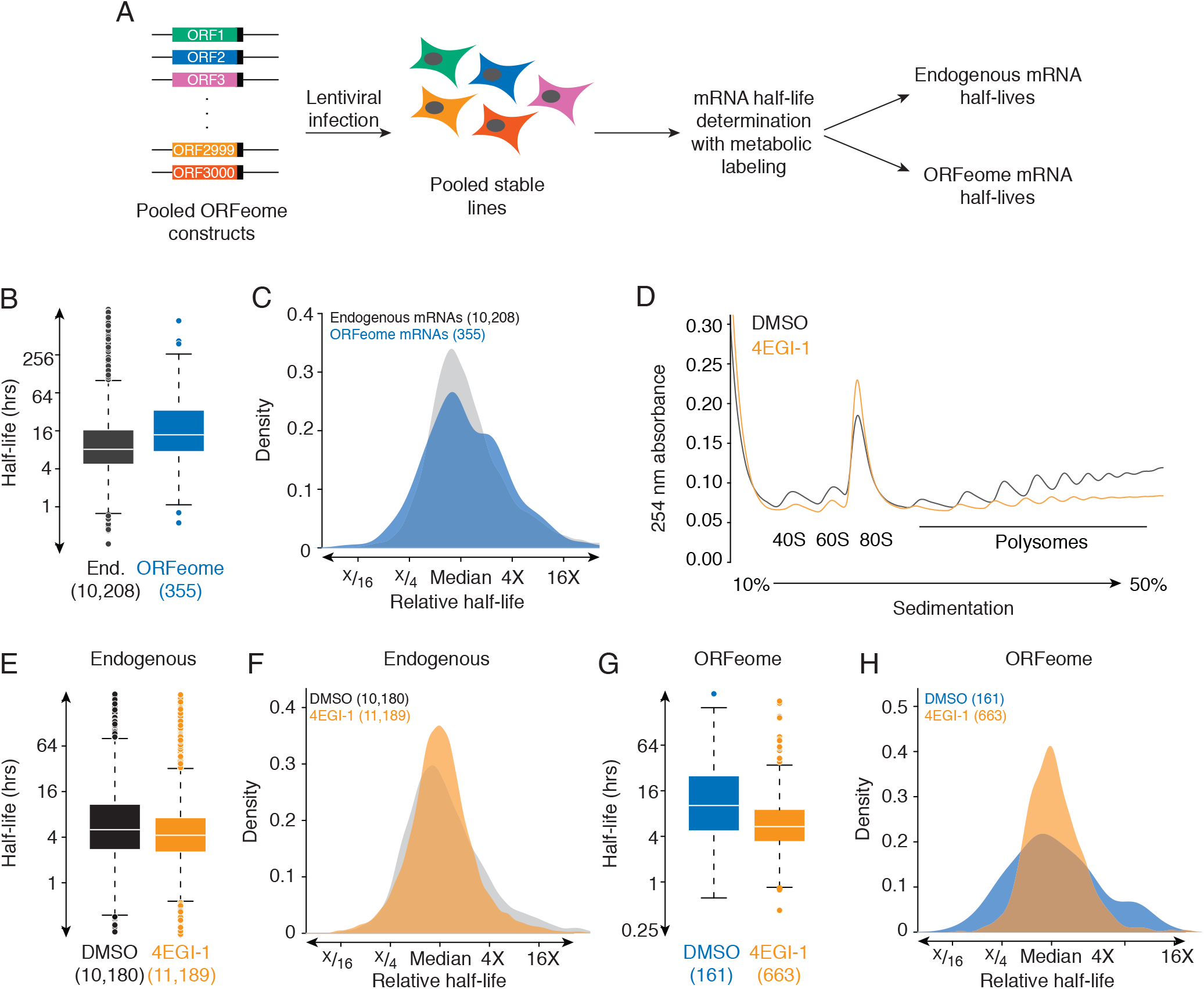
Coding sequences impact mRNA stability in human cells. (A) Schematic of the ORFeome workflow. The ORFeome collection contains ~16,000 full-length coding sequences corresponding to 14,000 genes in a lentiviral background. Each ORF derived from the ORFeome is flanked by invariant UTRs, and also contains a C-terminal V5 tag. Lentiviral pools containing ~3000 ORFeome clones were used to infect HEK293T and generate stable cell lines. Metabolic labeling was used to determine stabilities of endogenous and ORFeome-derived mRNAs in these stable lines. (B) Changing coding regions changes mRNA stability. Plotted are boxplots of half-lives for endogenous HEK293T (End.) and ORFeome-derived mRNAs (in blue and gray, respectively). The line represents the median half-life, and the box, 1^st^ and 3^rd^ quartiles. (C) ORFeome mRNAs show as much variation in stability as endogenous mRNAs. Plotted are density destributions of median-centered half-lives for endogenous HEK293T and ORFeome-derived mRNAs (in blue and gray, respectively). (D) Treatment with 4EGI-1 inhibits translation. Shown are A_254_ traces from sucrose density gradients of cell lysates from HEK293T cells treated with the translation inhibitor 4EGI-1 (orange) or DMSO (grey). (E) Transaltion inhibition destabilizes endogenous mRNAs. Plotted are boxplots of half-lives for endogenous HEK293T mRNAs with DMSO or 4EGI-1 treatment (in grey and orange, respectively). The line represents the median half-life, and the box, 1^st^ and 3^rd^ quartiles. (F) Translation inhibition has a minor effect on the variation in stability for endogenous mRNAs. Plotted are density destributions of median-centered half-lives for endogenous HEK293T in cells treated with DMSO or 4EGI-1 (in gray and orange, respectively). (G) Inhibition of translation destabilizes ORFeome-derived mRNAs and reduces the variation in stability. As in E, except for ORFeome mRNAs. DMSO, in blue; 4EGI-1, in orange. (H) Translation inhibition reduces the variation in stability for ORFeome-derived mRNAs. As in F, except for ORFeome mRNAs. DMSO, in blue; 4EGI-1, in orange. See also Figure S2 and Table S2.

We compared the distribution of half-lives from endogenous and ORFeome mRNAs (Figure 2B, C). ORFeome mRNAs were significantly more stable than endogenous mRNAs (*p* < 10^−15^), presumably due to the WPRE stabilizing element included in the invariant ORFeome 3’UTR (Zufferey et al., 1999). More strikingly, ORFeome mRNAs had as much, if not slightly more, variation in their stability as endogenous mRNAs (Figure 2C). Thus, we conclude that the ability of coding sequences to impact mRNA stability is a general phenomenon in human cells. Moreover, despite varying only in coding sequences and lacking any differences in UTR-based regulation, ORFeome mRNAs displayed the same range of stabilities as endogenous mRNAs, highlighting the importance of the ORF for regulating decay.

### Coding region determinants influence human mRNA stability only when translated

Given the known link between ORF-mediated regulation and translation, we asked whether the variation seen in ORFeome mRNA stabilities depended on translation. We treated cells with DMSO or 4EGI-1, a translation initiation inhibitor (Moerke et al., 2007), and again measured mRNA stabilites for endogenous and ORFeome-derived mRNAs. Using polysome profiling (Figure 2D) and incorporation of puromycin into nascent peptides (Figure S2D) (Schmidt et al., 2009), we confirmed that translation was broadly, although not completely, inhibited.

4EGI-1 had broad effects on transcript stability both for endogenous and ORFeome mRNAs. For endogenous genes, stabilities were poorly correlated between the DMSO and 4EGI-1 treatments (*r_s_* = 0.34, *p* < 10^−15^, Figure S2E, F). Endogenous mRNAs were also significantly destabilized in the presence of 4EGI-1 (*p* < 10^−15^, Figure 2E). Surprisingly, the variation seen in endogenous mRNA stability was also significantly, albeit modestly, reduced, (σ^2^ 4EGI-1/ σ^2^ DMSO = 0.65, *p* < 10^−15^, Figure 2F). These differences were more pronounced with the ORFeome mRNAs. ORFeome-derived transcripts were also less stable with 4EGI-1 treatment (*p* < 10^−10^, Figure 2G), but even more striking was the reduced variation in stability upon translation inhibition (σ^2^ 4EGI-1/ σ^2^ DMSO = 0.40, *p* < 10^−15^, Figure 2H). Taken together, we conclude that the impact of coding sequence on mRNA stability depends predominantly on translation.

### Coding regions influence mRNA stability independent of length, structure, and classical RBP-mediated regulation

We next wanted to determine what features in coding sequences affected mRNA stability. First, we examined three features previously linked with mRNA stability in eukaryotes: ORF length, secondary structure, and binding sites for RBPs (Duan et al., 2013; Geisberg et al., 2014; Neymotin et al., 2016; Schnall-Levin et al., 2011). Consistent with previous observations in many different eukaryotes, we observed a negative correlation between ORF length and mRNA stability for endogenous mRNAs (*r_s_* = –0.14, *p* < 10^−15^, Figure S3A). However, this relationship did not exist for ORFeome mRNAs (*r_s_* = 0.01, *p* = 0.9, Figure S3B), indicating that coding sequence length does not directly impact mRNA stability. The relationship between ORF length and stability observed with endogenous mRNAs could potentially be related by the underlying tendency for longer ORFs to also have longer 3’UTRs (*r_s_* = 0.18, *p* < 10^−15^), which themselves are negatively correlated with stability (*r_s_* = –0.15, *p* < 10^−15^)—rather than longer ORFs (or longer mRNAs) directly decreasing stability. These results also highlight the power of our ORFeome approach to separate correlative from causative effects on mRNA stability.

We next investigated the role of local secondary structure on mRNA stability. To do so, for each ORF, we calculated the folding energy in 100 bp sliding windows and measured the correlation between the minimum value and mRNA stability. For endogenous mRNAs, we found a weak relationship between local structure and mRNA stability (*r_s_* = 0.05, *p* < 10^−5^, Figure S3C); as with ORF length, this correlation failed to be significant for ORFeome-derived mRNAs (*r_s_* = 0.08, *p* = 0.1, Figure S3D). Thus, although extensive secondary structure within the coding sequence may destabilize individual transcripts, it does not provide a general explanation for the observed variation in ORFeome mRNA stability.

We next asked whether these differences could be explained by RBP or microRNA (miRNA)-mediated regulation. For each ORFeome-derived mRNA, we determined whether the coding region contained sites to the top five expressed miRNA families (Nam et al., 2014). Consistent with individual miRNA sites in the coding region having little impact on transcript stability (Grimson et al., 2007), half-lives for these site-containing ORFeome mRNAs were not significantly different from non-site mRNAs (*p* = 0.49, Figure S3E). Similarly, for Pumilio recognition elements, only six ORFs contained sites, and these half-lives did not differ from non-site ORFs. For AU-rich elements, we also did not observe any significant differences in transcript stabilities between ORFs that contained AU-rich elements and ORFs that did not contain AU-rich elements (*p* = 0.88, Figure S3F). Thus, we conclude that classical UTR-based regulation cannot explain differences in stability mediated by different coding sequences.

This conclusion is consistent with previous reports showing that miRNA sites in the ORF are rarely effective, likely because of the translating ribosome (Grimson et al., 2007; Gu et al., 2009). In support of this model, when translation was inhibited, AU-rich element-containing ORFeome mRNAs were now less stable than their no-site counterparts (*p* = 0.03, Figure S3G). This result also suggests that the some of the residual variation in ORFeome mRNA stability upon 4EGI-1 treatment (Figure 2H) may be due to RBP-based regulation that now has an opportunity to impact stability.

### Codon content is a major determinant of mRNA degradation in human cells

Because changing codon content altered mRNA stability in our reporters (Figure 1), we next explored the contribution of codon usage to differential ORF-mediated stability. To do so, for each non-stop triplet, we calculated the Spearman correlation between the frequency of the codon in each ORF and the associated half-life (so-called “codon stability coefficients” or CSCs), as we have done previously (Presnyak et al., 2015). We performed this analysis for endogenous and ORFeome-derived mRNAs in HEK293T cells, as well as in an additional dataset from endogenous mRNAs in HeLa cells (Figure 3A).

**Figure 3.**
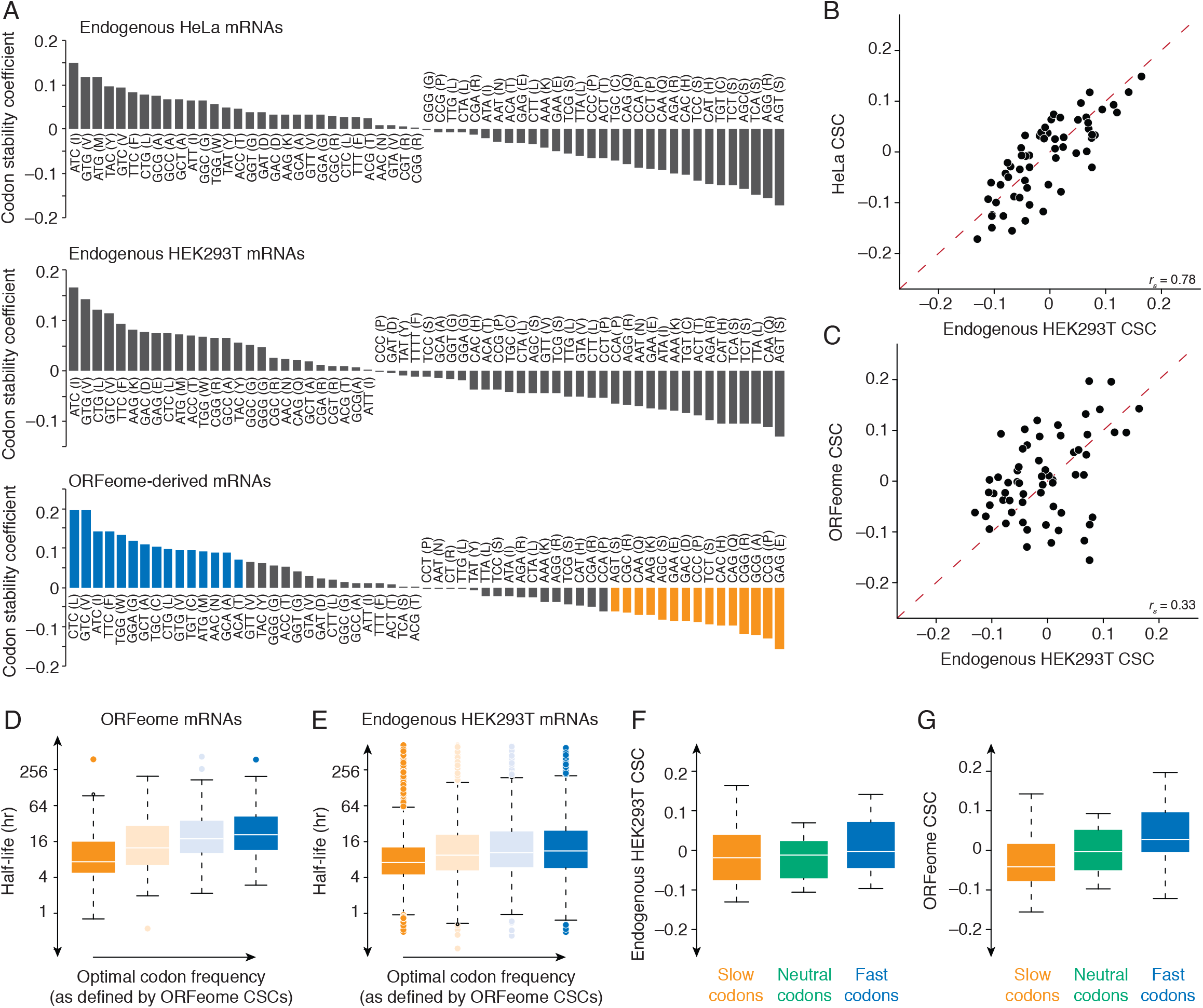
Codon usage is a determinant of mRNA stability in human cells. (A) Codons are differentially associated with stability. Shown are spearman correlations, for each codon, of their frequency with mRNA stability (codon stability coefficient; CSC) for endogenous HeLa mRNAs, endogenous HEK293T mRNAs, and ORFeome mRNAs. The 15 codons most associated with stability (as defined by the ORFeome collection) were designated as “optimal” (blue), while 15 codons most associated with instability were designated as “non-optimal” (orange). (B) HeLa and HEK293T cells have similar codon stability coefficients (CSCs). Plotted are the CSC values for endogenous HEK293T mRNAs compared to endogenous HeLa mRNAs (C) As in B, except comparing endogenous HEK293T and ORFeome-derived CSC values. (D) ORFeome mRNAs with more optimal codons are more stable. Shown are boxplots of mRNA half-lives for ORFeome mRNAs, binned into quartiles by the frequency of optimal codons. The line represents the median half-life, and the box, 1^st^ and 3^rd^ quartiles. (E) As in D, except for endogenous HEK293T mRNAs. (F) Endogenous HEK293T CSCs weakly correspond with pause scores. Using HeLa ribosome profiling, pause scores were calculated for each codon in the A site, and then codons were divided into three groups (slow in orange; neutral in green; fast in blue). Shown are boxplots for the corresponding CSC values as determined by endogenous HEK293T mRNAs. (G) As in F, except for ORFeome-derived CSCs. See also Figures S3 and S4.

Some codons behaved similarly. For instance, ATC was associated with stability in all three datasets. However, others, such as GAG, were not: in the ORFeome analysis, it was the most instability-associated codon, but in the endogenous HEK293T analysis, it was associated with stability. Across all 61 codons, the CSC values derived from endogenous mRNAs in HeLa and HEK293T cells were remarkably similar (*r_s_* = 0.78, *p* < 10^−15^, Figure 3B); in fact, the CSC values were more similar than the underlying half-lives themselves (*p* < 10^−4^, Fisher’s r-to-z transformation,). In contrast, although CSC values for the endogenous and ORFeome analyses were significantly correlated (*r_s_* = 0.33, *p* = 0.009; Figure 3C), this relationship was weaker than that between HEK293T and HeLa cells (*p* = 0.0002, Fisher’s r-to-z transformation). One possibility is that some of the similarity between HEK293T and HeLa values can be explained by similar ORF- and UTR-mediated regulation, and that the reduced correlation with the ORFeome values reflects the lack of UTR-mediated regulation for ORFeome mRNAs.

We next took the fifteen codons most associated with stability and instability in our ORFeome analysis and considered these as stable and unstable codons, respectively. As expected, half-lives for ORFeome mRNAs were significantly associated with the combined frequency of the stabilizing codons (*r_s_* = 0.36, *p* < 10^−11^, Figure 3D). Similarly, half-lives for endogenous HEK293T mRNAs and endogenous HeLa mRNAs were significantly correlated with this combined frequency, although to a lesser extent (HEK293T *r_s_* = 0.18, *p* < 10^−15^, Figure 3E; HeLa *r_s_* = 0.17, *p* < 10^−15^, Figure S4A). In contrast, when we performed computational frame-shift controls, the correlation between ORFeome CSC values and endogenous HEK293T mRNA stabilities was lost (*r_s_* = 0.07 and 0.01). Together, these data are consistent with a model where different codons rely upon translation to impact mRNA stability, rather than the sequences, in and of themselves, mediating different decay rates.

### Instability-associated codons are translated more slowly

Based on evidence from yeast (Hanson et al., 2018), the current view is that the elongating ribosome takes longer to move through some codons than others, and that these slowly elongating codons stimulate mRNA decay. Given this model, we next investigated the relationship between ribosome elongation speed and our three sets of CSC values. To determine relative elongation speed, we analyzed ribosome profiling datasets derived from HeLa cells (Arango et al., 2018). After inferring the A-site codon identity for each fragment, we calculated the relative frequency of each codon in the A site, reasoning that slower codons will have more A-site associated reads. There was only a weak relationship between relative elongation speed and endogenous CSC values (HEK293T: *r_s_* = −0.14, *p* = 0.3; HeLa: *r_s_* = –0.22, *p* = 0.9; Figure 3F, Figure S4B). Strikingly, this relationship was substantially stronger and significant for ORFeome-derived CSC values (*r_s_* = –0.36, *p* = 0.004, Figure 3G), likely because of the lack of confounding UTR-mediated regulation. Thus, we conclude that instability-associated codons are translated more slowly by the elongating ribosome.

### Amino acid identity is also a major determinant of mRNA stability in human cells

In yeast and other organisms (Presnyak et al., 2015), the impact of different codons can be explained by differences in tRNA abundance. We thus compared relative tRNA abundance in HEK293T cells (Zheng et al., 2015) with our codon stability metrics (Figure 4A). As expected, tRNA abundance did correlate with endogenous HEK293T CSC values (*r_s_* = 0.31, *p* = 0.02). Importantly, this correlation is not nearly as robust as that seen in yeast (Presnyak et al., 2015); thus, other aspects of codon identity could be at play. Consistently, tRNA concentration effects on mRNA decay are weakened in the ORFeome CSC values (*r_s_* = 0.05, *p* = 0.7). Further, there was only a weak relationship between ribosome elongation speed and tRNA abundance in general (*r_s_* = 0.10, *p* = 0.5; Figure 4B).

**Figure 4.**
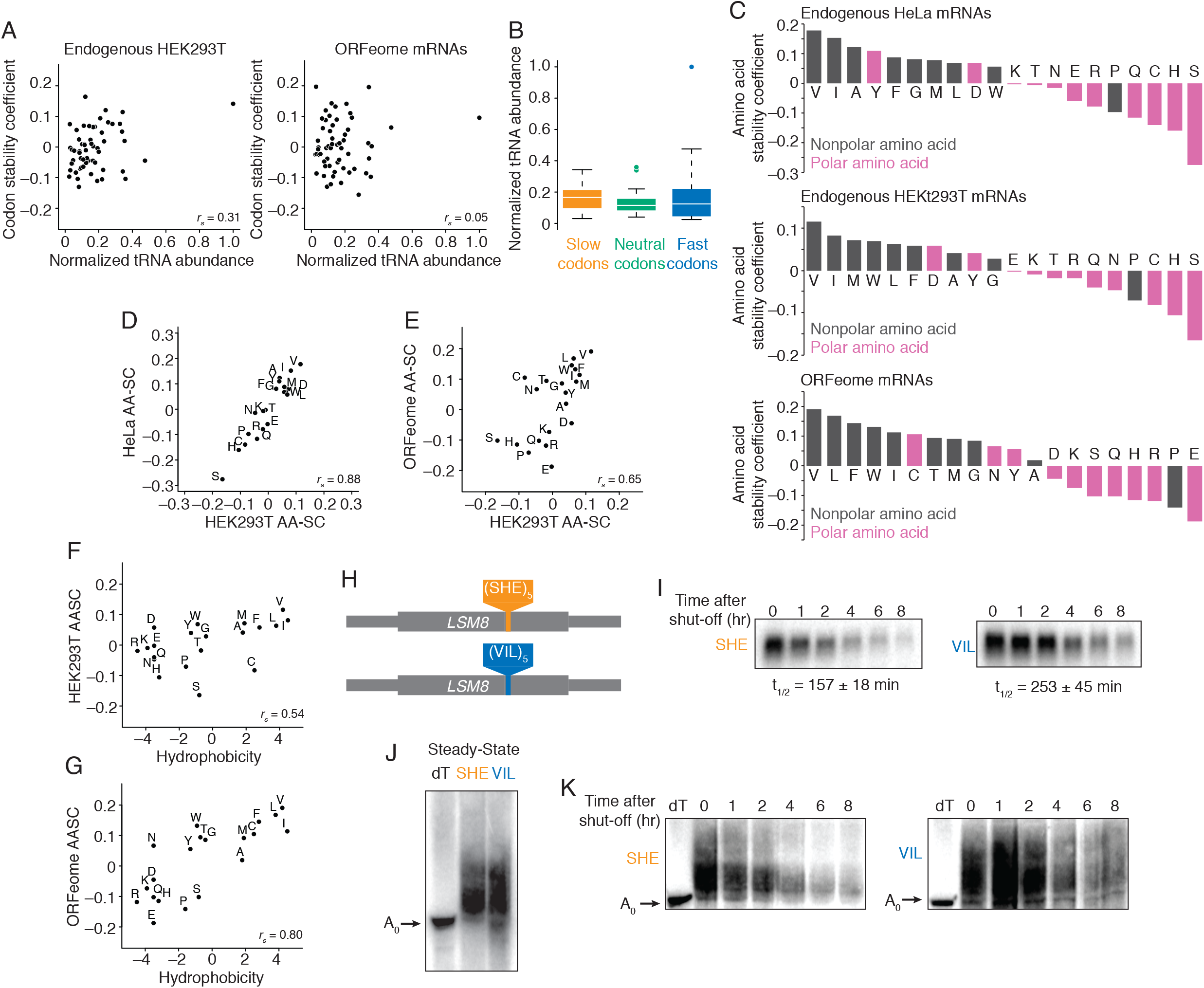
Amino acid usage is a major determinant of mRNA stability in human cells. (A) Codon stability coefficients (CSCs) have little relationship to tRNA abundance. Plotted are the normalized tRNA abundance in comparison to CSC values for endogenous HEK293T and and ORFeome mRNAs (left and right panels, respectively). (B) tRNA abundance has little impact on elongation speed. Codons were divided into thirds by their A-site pause scores (slow in orange; neutral, green; fast, blue). Shown are boxplots for the abundance of corresponding tRNAs. The line represents the median half-life, and the box, 1^st^ and 3^rd^ quartiles. (C) Amino acids are differentially associated with stability. Shown are spearman correlations, for each amino acid, of their frequency with mRNA stability (amino acid stability coefficient or AASC) for endogenous HeLa mRNAs, endogenous HEK293T mRNAs, and ORFeome mRNAs. Polar amino acids (in pink) have charged or highly electronegative side chains; nonpolar amino acids (dark gray) have aliphatic and weakly electronegative side chains. (D) HeLa and HEK293T have similar AASCs. Plotted are the AASC values for endogenous HEK293T mRNAs compared to endogenous HeLa mRNAs (E) As in B, except comparing endogenous HEK293T and ORFeome-derived AASC values. (F) Hydrophobic amino acids are associated with stability. Plotted are the hydrophobicity scores (Kyte and Doolittle, 1982) for each amino acid compared to their stability coefficient for endogenous HEK293T mRNAs. (G) As in F, except for ORFeome-derived AASC values. (H) Schematic diagram of *LSM8* reporter constructs. In the middle of the *LSM8* coding region, five repeats of instability-associated amino acids (S, H and E) or stability-associated amino acids (V, I, and L) were inserted. (I) Insertion of instability-associated amino acids destabilizes the *LSM8* reporter mRNA. Transcriptional shut-off experiments were performed in Tet-Off HEK293 cells, and *LSM8* mRNA levels were determined by northern blots. Timepoints correspond to the time after the addition of doxycycline. t½ corresponds to the half-life (min) ± standard deviation (n=4). See Figure S5C for loading control. (J) The destabilized *LSM8* reporter mRNA has shorter poly(A) tails. High resolution northern blotting was performed to measure poly(A)-tail lengths on the SHE and VIL *LSM8* mRNAs. Arrow indicates deadenylated mRNA species; dT, oligo(dT)/RNase H treated mRNA control. (K) *LSM8* reporter mRNAs are deadenylated. Transcription of the SHE and VIL *LSM8* reporters was shut-off, as in I, and poly(A)-tail lengths were measured by high-resolution northern blotting. Timepoints represent time elapsed after transcription shutoff with 2 μg/mL doxycycline. Arrow indicates deadenylated mRNA species; dT, oligo(dT)/RNase H treated mRNA control. See also Figure S5 and Tables S1 and S2.

We hypothesized that something other than tRNA abundance could explain the observed codon-specific differences in elongation speed and impact on stability. In analyzing our CSC values, we also noted that codons encoding the same amino acids tended to behave similarly, leading us to investigate the impact of amino acid composition on mRNA stability. Analogous to our CSC values, we determined an amino acid stabilization coefficient (AASC) for each amino acid in our three half-life datasets (Figure 4C). As before, the amino acid metrics were very similar between HeLa and HEK293T cells (*r_s_* = 0.88, *p* < 10^-15^, Figure 4D) and between endogenous and ORFeome-derived mRNAs (*r_s_* = 0.65, *p* = 0.002, Figure 4E).

Strikingly, in all three datasets, hydrophobic amino acids were associated with stability, while polar and charged amino acids with instability (Figure 4F, G) (Kyte and Doolittle, 1982). In fact, AASC values from endogenous HEK293T mRNAs were significantly associated with hydrophobicity (*r_s_* = 0.54, *p* = 0.02, Figure 4F)—a relationship that was even stronger for the ORFeome-derived values (*r_s_* = 0.80, *p* < 10^−4^, Figure 4G). For example, those coding sequences containing more valine residues were more stable than those with fewer valines in all three half-life datasets (endogenous HEK293T mRNAs: *r_s_* = 0.12, *p* < 10^−15^; HeLa mRNAs: *r_s_* = 0.18, *p* < 10^−15^; ORFeome mRNAs: *r_s_* = 0.19, *p* < 10^−3^; Figure S5A). One notable exception was proline, which was associated with instability in all three datasets, despite being nonpolar.

Nonetheless, there were some differences between AASC values derived from ORFeome mRNAs and endogenous mRNAs. For instance, serine was most highly associated with instability in endogenous mRNAs (HEK293T: *r_s_* = –0. 16, *p* < 10^−15^; HeLa: *r_s_* = –0.28, *p* < 10^-15^), but it was more weakly associated with instability in the ORFeome transcripts (*r_s_* = –0.10, *p* = 0.06; Figure S5B). In contrast, glutamate was very strongly associated with instability in the ORFeome mRNAs (*r_s_* = –0.19, *p* < 10^−3^), and yet its use had little association with instability for either of the two endogenous mRNA sets (HEK293T: *r_s_* = 0.00, *p* = 0.8; HeLa: *r_s_* = –0.06, *p* < 10^−10^).

To determine whether these relationships were causative, we engineered reporters containing the ORF sequence for a relatively small gene [*LSM8* (291 nt, 96 amino acids)] with a mid-ORF insertion of 15 amino acids (45 nt) (Figure 4H). These amino acid stretches were composed of 5 tandem repeats of either “stabilizing” amino acids (valine, isoleucine, and leucine) or “destabilizing” amino acids (serine, histidine, and glutamate) based on the most extreme AASC values from all three datasets. Importantly, constituent codons were randomly assorted to ensure approximately equal codon usage and reduce local GC content extrema (Table S1). We then performed single-gene transcriptional shutoff using the Tet-Off system and determined mRNA half-lives by northern blot. The reporter containing the destabilizing amino acid insertion was significantly less stable than that with the stabilizing amino acids (Figure 4I, Figure S5C; *p* = 0.007, two-tailed t-test), thus indicating that the relationship between amino acid content and mRNA stability is causative.

In yeast, non-optimal codons recruit the decay machinery, leading to increased deadenylation (Radhakrishnan et al., 2016; Webster et al., 2018). We next asked whether there might be a similar mechanism at work in human cells. Using high-resolution northern blots, we determined poly(A)-tail lengths on our stable and unstable *LSM8* reporter mRNAs. The unstable *LSM8* reporter mRNA showed shorter overall steady-state tail length than the stable reporter (Figure 4J), and a higher proportion of transcripts were deadenylated over the course of transcriptional shut-off experiments (Figure 4K). Taken together, these data strongly support the model that primary protein sequence influences mRNA half-life in human cells, and that it likely does so through a deadenylation-based mechanism.

### Amino acid identity influences elongation speed

Given that amino acids like proline and glutamate are associated with instability and are also known to slow elongation (Artieri and Fraser, 2014; Gardin et al., 2014), we wondered whether this trend held more generally. Using a similar analysis to our codon elongation rate metric, we calculated relative pause scores for each amino acid (when in the inferred A site). Consistent with a model where slow elongation is associated with instability, stabilizing amino acids (such as valine and leucine) showed little evidence of pausing, while destabilizing amino acids (like glutamate and histidine) had the strongest pausing signal (Figure 5A).

**Figure 5.**
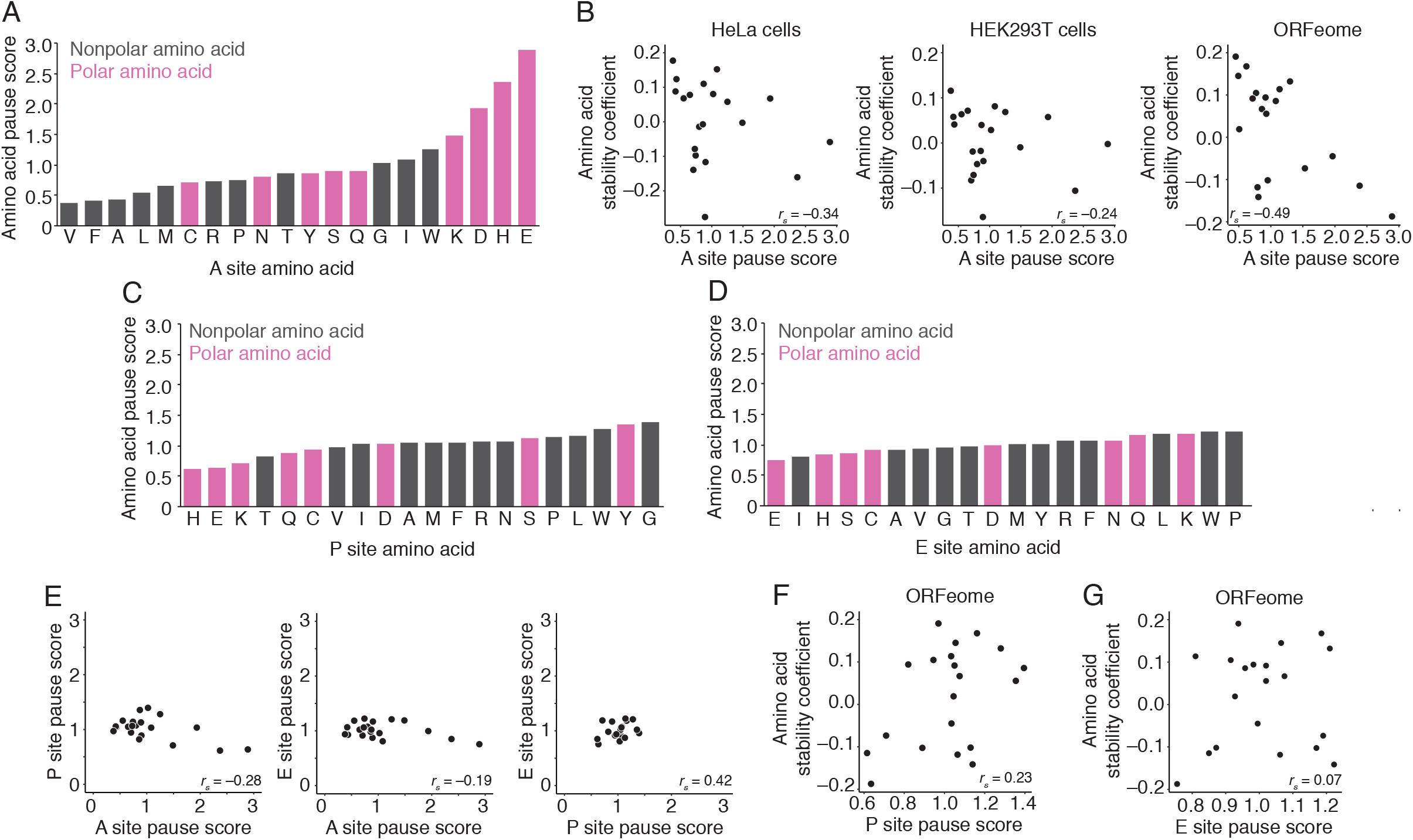
Instability-associated amino acids are translated slowly. (A) Amino acids, when in the A-site, are translated at different rates. Plotted are the pause scores for each amino acid when in the predicted A-site (see methods for details). Nonpolar amino acids, grey; polar amino acids, pink. (B) Amino acid stability coefficients correlate with A-site pause scores. Shown are plots comparing A-site pause scores for each amino acid with its stability coefficient, as defined by endogenous HeLa, endogenous HEK293T, and ORFeome mRNAs (left, middle, and right, respectively). (C) As in A, except for the P site. (D) As in A, except for the E site. (E) A-site pause scores correlate poorly with P- and E-site pause scores. Plotted are the pairwise comparisons for A-, P-, and E-site pause scores. (F) ORFeome amino acid stability coefficients poorly correlate with P-site pause scores. Shown are plots comparing P-site pause scores for each amino acid with its stability coefficient, as defined by ORFeome mRNAs. (G) As in F, except for E-site pause scores.

Although the A-site pause scores showed only a weak association with both sets of endogenous AASC values (HEK293T: *r_s_* = –0.24, *p* = 0.3; HeLa: *r_s_* = –0.34, *p* = 0.1; Figure 5B), there was a significant negative correlation between ribosome pausing and ORFeome-derived stability values (*r_s_* = –0.50, *p* = 0.03; Figure 5B). As with our codon analysis, this relationship between pausing and instability values was likely stronger for the ORFeome values because these lack confounding effects of the UTRs. One interesting exception was proline, which had a relatively weak A-site pause score, but was strongly associated with instability in all three datasets. Proline is both a poor amino acid bound donor and acceptor (Wohlgemuth et al., 2008), and so it is likely that A-site pause scores do not completely capture its contribution to reducing elongation speed. Importantly, these observations only held true for amino acid content in the A site. As opposed to A-site pause scores, there was little variation between amino acids for P- and E-site pause scores, and we observed no correlation between ORFeome AASC and ribosome pausing for neither the P-nor E-site amino acid (Figure 5C-G). Taken together, we conclude that in humans, amino acid content within the ribosomal A site influences elongation; like the ribosome pausing trigged by tRNA limitations, relative elongation rate over amino acids is sensed by the mRNA degradation machinery.

### The relative contribution of amino acid and codon use to mRNA stability varies in different species

Codon effects are inherently entwined with amino acid effects. We asked, therefore, to what extent differences between amino acids (as opposed to differences between synonymous codons) contributed towards our observations on transcript stability. We first focused on *S. cerevisiae*, where elongation speed is known to be predominantly determined by the decoding step and tRNA concentration (Hanson et al., 2018). We binned synonymous codons and plotted their CSC values (Figure 6A). Consistent with previous work (Hanson et al., 2018; Presnyak et al., 2015), there was wide variation in CSC values for synonymous codons, and these values reflected tRNA abundance (Figure 6A). In HeLa cells, however, less variation between synonymous codons was observed; instead, CSC was driven predominately by amino acid identity (Figure 6B).

**Figure 6.**
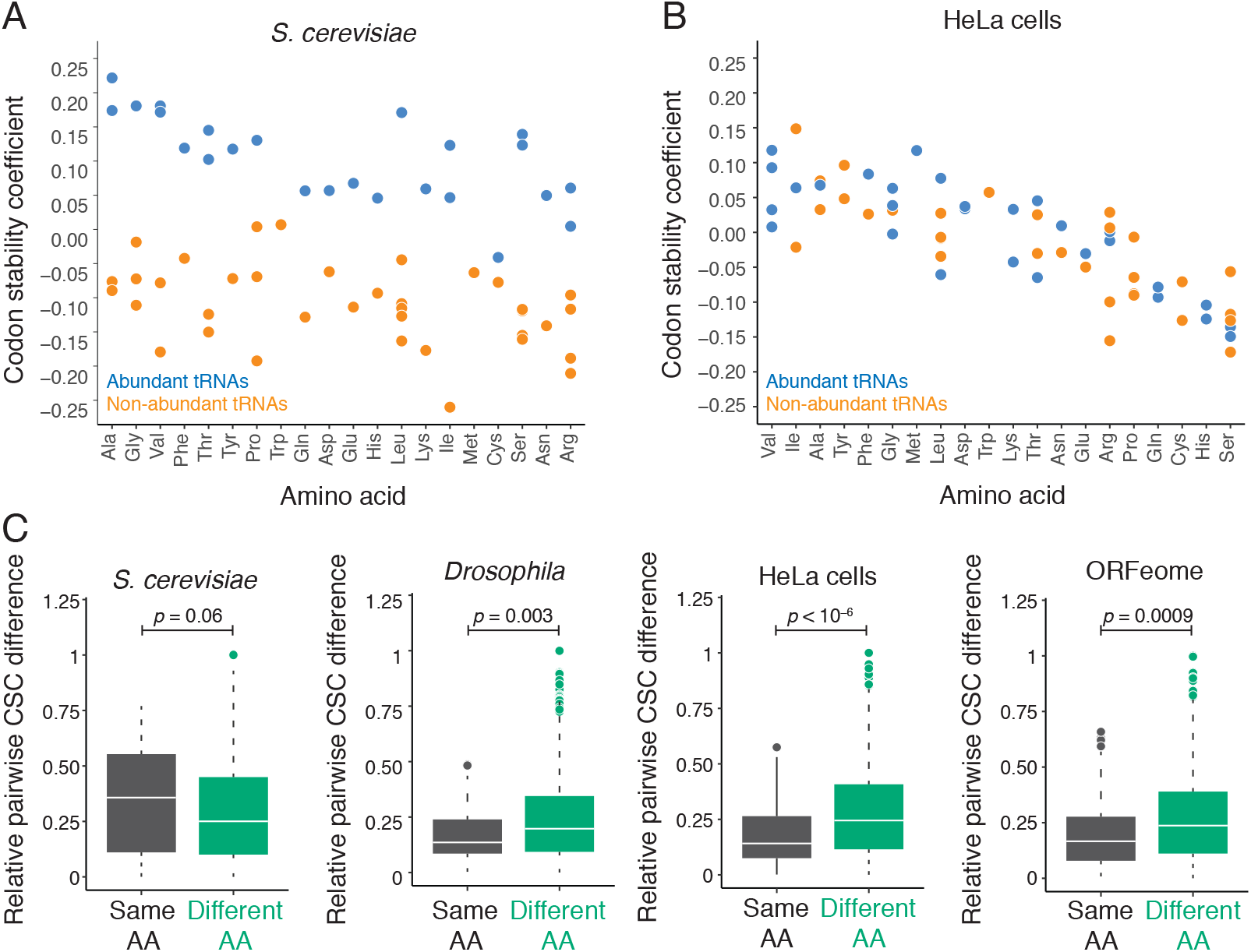
Amino acid usage is a more potent regulator than codon usage in control of human mRNA stability. (A) Comparison of codon and amino acid effects on stability in *S. cereviaise*. Codons were grouped by their encoded amino acid, and codon stability coefficient (CSC) values are plotted. Those decoded by high-abundance tRNAs are in blue, and those by low-abundance tRNAs are in orange. (B) As in A, except for HeLa cells. (C) In metazoans, differences between codons are predominantly determined by differences between amino acids. For each pair of codons, the absolute difference in the corresponding CSC values was calculated and then normalized to the maximal difference (to correct for differences in overall variance between organisms). Pairs of codons were binned into these encoding the same or different amino acid (n = 87, in grey, and n = 1742, in green, respectively). Shown are boxplots of those differences for *S. cerevisiae*, *Drosophila*, HeLa, and ORFeome-derived mRNAs. Significance determined by Wilcoxon rank sum test. See also Table S3.

To quantitatively asses the relative contribution of codon vs. amino acid identity, we developed a metric that determines the relative impact of synonymous and non-synonymous codons on mRNA stability. Here, we took each pair of synonymous and non-synonymous codons and calculated the differential between their CSC values. This score was then normalized to the largest score across all codon combinations for each organism. In *S. cereviaise*, consistent with the importance of tRNA abundance rather than amino acid identity, synonymous codon usage gave a large range in observed CSCs (Figure 6C). For *S. pombe* and trypanosomes, a similar pattern was observed (Figure S6). These data indicate that for these organisms, synonymous codon usage, rather than amino acid identity, correlate best with differences in mRNA stability. However, in metazoans (such as *Drosophila*, zebrafish, and human cells), there was more variation between non-synonymous codons than synonymous codons (Figure 6C, Figure S6), indicating that amino acid effects dominate over codon effects. Thus, while tRNA abundance appears to be a dominant feature in dictating mRNA stability in fungi and protists, in metazoans it is the encoded amino acid. These data hint that translational elongation rate is differentially impacted by distinct rate-limiting events throughout evolution. Nonetheless, any perturbation that slows elongation has the same net influence on mRNA stability.

### Amino acid content coordinates mRNA stability within gene families

Previous studies in yeast demonstrated that expression of functionally related genes can be orchestrated at the level of mRNA stability and that much of this phenomenon occurs through coordinated synonymous codon use (Presnyak et al., 2015; Wang et al., 2002). In human cell lines, we observed a similar phenomenon where functionally related gene groups have similar mRNA half-lives (Figure 7A). For instance, in HeLa cells, key metabolic genes, such as TCA cycle enzymes, were highly stable (median half-life: 8.53 hrs), while regulatory proteins, such as Serine/Arginine-rich splicing factors (SRSF), were highly unstable (median half-life: 2.83 hrs). Given that amino acid identity is a major driver of mRNA stabilities in human cells, coordinate gene expression at the mRNA level could, in theory, be driven by primary amino acid sequence.

**Figure 7.**
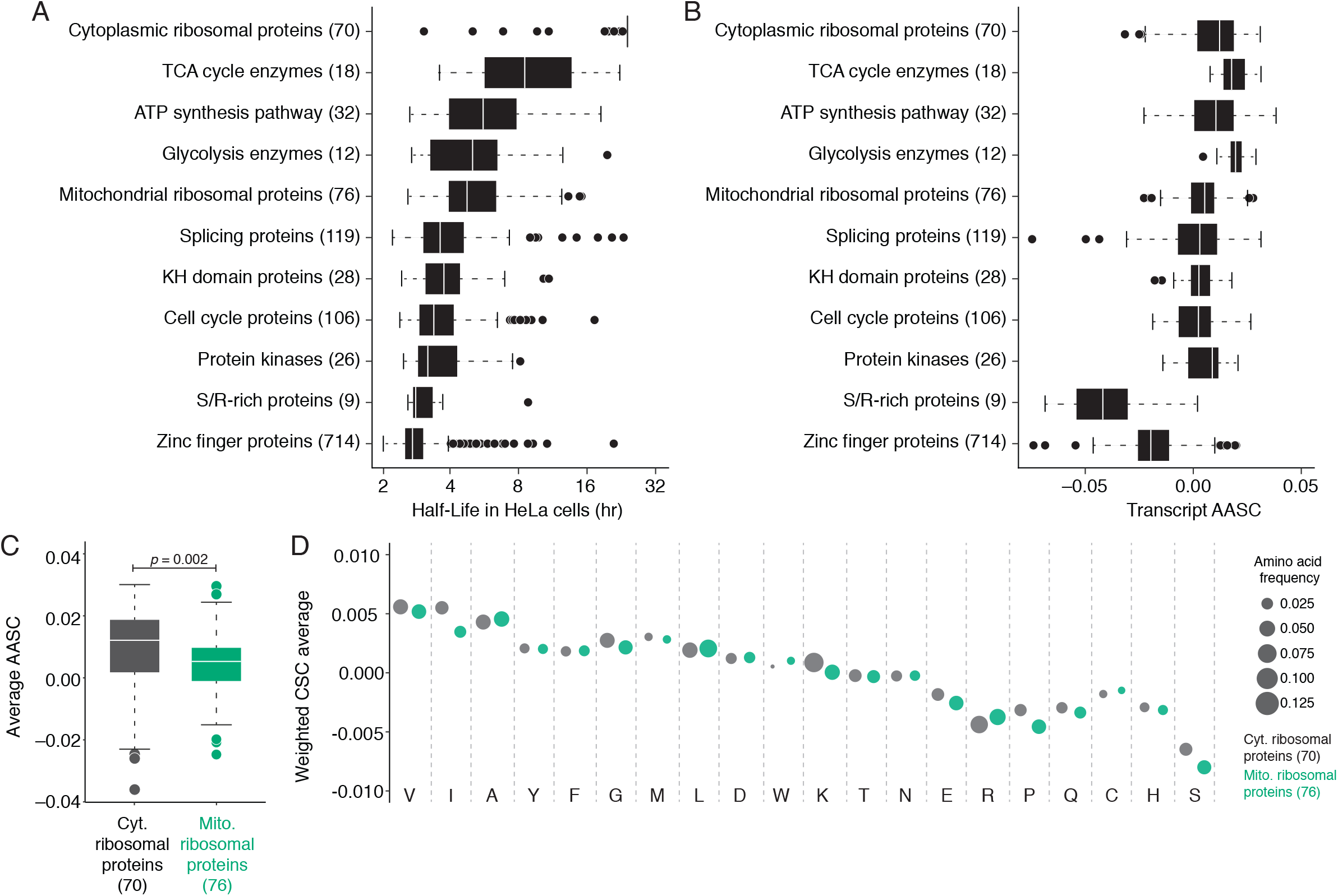
Similar amino acid content helps coordinate control of mRNA stability among functionally related genes. (A) Gene groups show similar half-lives. Shown are boxplots of mRNA half-lives in HeLa cells for the indicated gene groups. (B) Gene groups show similar average AASCs. Shown are boxplots of AASC values in HeLa cells for the indicated gene groups. (C) Mitochondrial ribosomal protein genes have lower AASCs than cytoplasmic ribosomal protein genes. Shown are boxplots of AASC values for cytoplasmic and mitochondrial ribosomal protein genes (blue and red, respectively). Significance was determined by Wilcoxon rank sum test. (D) Mitochondrial ribosomal protein genes use more unstable amino acids and codons than cytoplasmic ribosomal protein genes. Each amino acid is represented by a dot, where its size corresponds to its frequency, and is plotted by the weighted CSC value. Shown are the distributions for cytoplasmic and mitochondrial ribosomal protein genes (in blue and red, respectively). See also Figure S6.

To explore potential relationships between mRNA half-life and amino acid content across gene families, we calculated average AASC across these same transcripts (Figure 7B). Highly stable gene groups showed higher average AASC (TCA cycle enzyme median AASC: 0.0178), while unstable gene groups showed markedly lower average AASC (SRSF median AASC: –0.0419), supporting a model where differences in amino acid composition can help coordinate mRNA half-lives within functionally-related gene groups.

We further examined differences in amino acid and codon usage between two groups of interest, cytoplasmic and mitochondrial ribosomal proteins. Despite being closely related, these two groups have significantly different half-lives (*p* = 0.0009, Wilcoxon test; Figure 7A). Reflecting differences in amino acid content, these two groups also differ significantly in AASC values (*p* = 0.002, Wilcoxon test; Figure 7C). Of note, mitochondrial ribosomal proteins were found to contain more destabilizing amino acids, such as serine and glutamate, while cytoplasmic ribosomal proteins contained more stabilizing amino acids, such as valine and isoleucine (Figure 7D). In addition, these two groups also showed differences in synonymous codon usage, such that mitochondrial ribosomal protein genes tended to contain codons with lower CSC values (Figure 7D). Taken together, these analyses demonstrate that both codon usage and amino acid content contribute to the observed differences in half-life between functionally related gene groups. Moreover, these data strongly suggest that alterations in amino acid content could be a powerful evolutionary driver of overall gene expression levels.

## DISCUSSION

Despite the elucidation of most major mechanisms of mRNA degradation over the past two decades, much about how mRNA stability is regulated in humans remains unknown. Here, we have demonstrated that translational elongation rate is an evolutionarily conserved determinant of mRNA stability. First, using reporter systems, we show that codon content impacts transcript stability. Reporters containing largely non-optimal codons were significantly less stable than transcripts containing largely optimal codons. These results are not isolated to a few, select mRNAs; rather, general correlations between codon usage, elongation speeds, and mRNA stability hold on a transcriptome-wide scale in humans, indicating that this regulatory mechanism is broadly conserved.

Importantly, the human ORFeome collection brought these effects into even more focus. Unlike endogenous genes, mRNAs expressed from the ORFeome collection contain the same UTRs, promoters, and poly(A) sites, and so regulation from elements outside the ORF (such as on translational initiation rates) are not in play for ORFeome-derived mRNAs. The relationship between codon effects, mRNA stability, and elongation rates was even stronger for these transcripts compared to endogenous genes, demonstrating a causative, translation-dependent relationship between coding elements and regulation of mRNA stability.

Codon effects on mRNA decay are driven by tRNA abundance in yeast (Presnyak et al., 2015). In humans, we also see that tRNA concentration can powerfully impact mRNA degradation rate; reporter constructs (Fig. 1) whose optimality is determined based on tRNA concentration exhibit effects on mRNA decay identical to that seen in yeast. Importantly, however, analyses in humans revealed that amino acid identity is also a major driver for mRNA decay rates, such that polar amino acids are generally destabilizing and nonpolar amino acids are stabilizing. The effect of the primary protein sequence on mRNA stability is not merely a correlative one. When we inserted a stretch of codons encoding polar or nonpolar amino acids into an *LSM8* reporter gene, the two mRNAs had different stabilities. Despite only varying in 15 codons, the variant encoding polar amino acids was more unstable and had a shorter poly(A) tail. Importantly, the codons for amino acids most associated with instability also showed the strongest ribosome pausing when in the A site. Together, our results lead to a model that amino acid identity and chemical properties impact ribosomal A-site incorporation and, in turn, overall translational elongation rate, and so encoded protein has a direct impact on the stability of its mRNA.

We hypothesize that the distinction between tRNA concentration as opposed to amino acid identity in controlling transcript stability result from subtle evolutionary changes in the rate-limiting steps for translational elongation. In fungi and protists, decoding appears to limit translational elongation, while amino acid effects modulate elongation in metazoans. The net effect on mRNA stability, however, is the same: slowing translation elongation results in mRNA destabilization, most likely by stimulating deadenylation.

A key question raised by this study is how amino acid identity within the A site changes elongation speeds. The observation that polar amino acids behave similarly raises the possibility that these amino acids have a chemical nature or physical geometry that is difficult for human ribosomes to either accommodate or move into the growing polypeptide chain, thereby creating a pause that can then be sensed by the mRNA decay machinery. Indeed, the ribosome is known to have difficulty with certain amino acids (Artieri and Fraser, 2014; Gardin et al., 2014). For instance, polyproline present within the growing polypeptide chain is strongly inhibitory to ribosome elongation (Wohlgemuth et al., 2008). Similarly, a polypeptide bond can be difficult to form between certain tRNA-amino acid combinations due to slight misalignments in the tRNA backbone and/or amino acid geometry. Indeed, the evolution of the translational elongation factor, eIF-5A, demonstrates this constant adaptive need for substrate alignment within the ribosome (Gutierrez et al., 2013; Saini et al., 2009). eIF-5A nudges the P-site tRNA, lining it up appropriately with the A-site tRNA to allow chemistry to occur (Melnikov et al., 2016). Although other possibilities (e.g., differences in tRNA charging) are also possible, we favor a model where polar amino acids in the A site creates subtle pauses in elongation due to inappropriate aligning with the peptidyl transferase center. These subtle, yet likely additive, pausing events are sensed by the mRNA degradation machinery to elicit transcript destruction.

### Protein sequence dictates mRNA expression

We and others observed that genes of similar physiological function coordinate their expression through coordinated mRNA stability (Herzog et al., 2017; Presnyak et al., 2015; Wang et al., 2002). In yeast, this coordination is achieved by common usage of synonymous codons within gene classes. Because each of the 61 codons is read at distinct rates (due to differences in tRNA concentration), codon composition has a strong impact on overall expression level through translational elongation rate and mRNA degradation rate (Figure 8). Across evolutionary time, selection for synonymous codon usage will modulate gene expression, but it does so without impacting protein amino acid composition.

**Figure 8.**
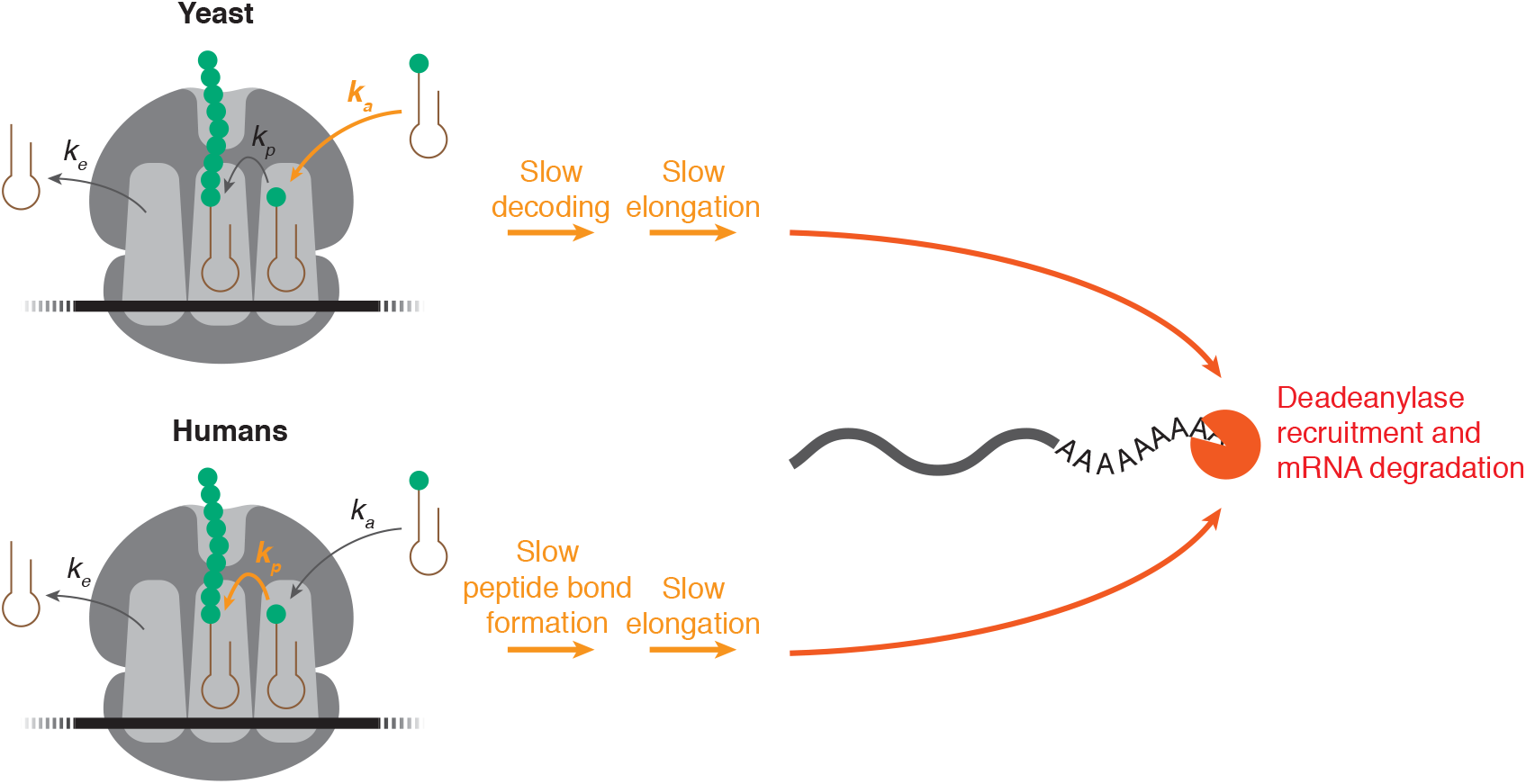
Model: mRNA stability is determined by speed of translation elongation. This schematic demonstrates a model for mRNA stability as a function of translation elongation speed. We propose that the combined rate of tRNA decoding/accommodation (*K_a_*), rate of peptidyl transfer from P-site tRNA to A-site tRNA (*K_p_*), and rate of deacylated tRNA exit from the E-site (*K_e_*) dictate speed of ribosome transit. Ribosome speed is sensed by yet-to-be-determined mechanisms and communicates with the mRNA decay machinery to determine mRNA decay rate. Here, “destabilizing” codons and amino acids can slow the rate of tRNA accommodation (*K_a_*) and the rate of peptidyl transfer (*K_p_*), leading to slowing of elongation and subsequent targeting for decay.

Here, however, we demonstrate that human mRNA stabilities are coordinated at the codon level predominantly through amino acid composition rather than tRNA abundance—in other words, the overall primary sequence of a polypeptide hardwires the turnover rate of its corresponding mRNA (Figure 8), and many protein sequences will restrict mRNA expression. For some genes, the intrinsic ability of the protein sequence to restrict mRNA expression may be a feature that evolution has taken advantage of in order to enable dynamic expression patterns. On the other hand, for a gene whose protein product needs to be highly expressed, this ORF-mediated regulation must be either overcome (*e.g*., by increased transcriptional buffering) or blunted by other post-transcriptional regulation (*e.g*., UTR-based mechanisms). The latter concept suggests that evolution of protein sequence could, in fact, be an adaptive response to the need for coordinated mRNA stability within some gene families, especially those subject to high translation initiation rates. Further studies will be required to understand the interplay between UTR-based regulation and ORF composition in determining mRNA half-life in higher eukaryotes.

## ACKNOWLEDGEMENTS

We thank Dr. Julie Claycomb, Dr. Jeff Kieft, and members of the Coller, Oberdoerffer, Ellis, and Rissland labs for thoughtful discussions. We thank Dr. Mikko Taipale and Dr. Jason Moffat for help with the ORFeome collection. Support was provided from NIH grants R35GM128680 (OSR) and R01GM118018, R01GM125086 (JC), T32GM0080056 (JC and MF) and from CIHR grant PJT-148746 (JE).

## AUTHOR CONTRIBUTIONS

Conceptualization, O.S.R and J.C.; Methodology, M.E.F., A.N., T.J.S, D.A., G.H., J.E., S.O., J.C., and O.S.R.; Data Analysis and Curation, M.E.F., A.N., T.J.S, D.A., G.H.; Investigation and Validation, M.E.F., A.N.,T.J.S, D.A., G.H.; Writing - Original Draft, O.S.R, and J.C.; Writing - Review & Editing, M.E.F, A.N., O.S.R., and J.C.; Supervision and Funding Acquisition, O.S.R., J.C., and J.E.

## SUPPLEMENTAL FIGURE LEGENDS

**Figure S1.**
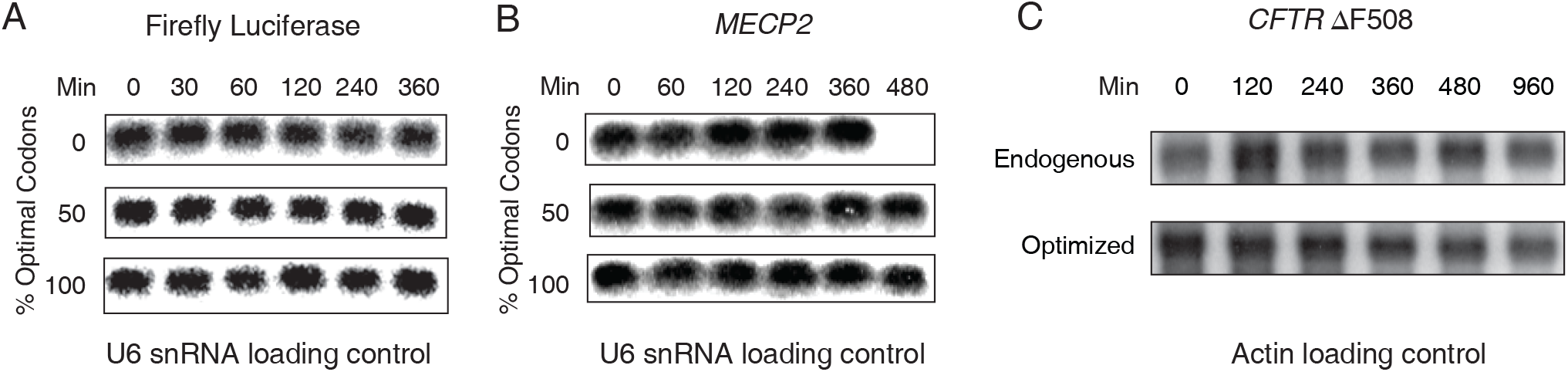
Optimal codon usage modulates mRNA stability in human cells, related to Figure 1. (A) U6 snRNA northern analysis for transcription shut-off experiments for the firefly luciferase variants shown in Figure 1C. (B) As in A, except for *MECP2* and Figure 1D.

**Figure S2.**
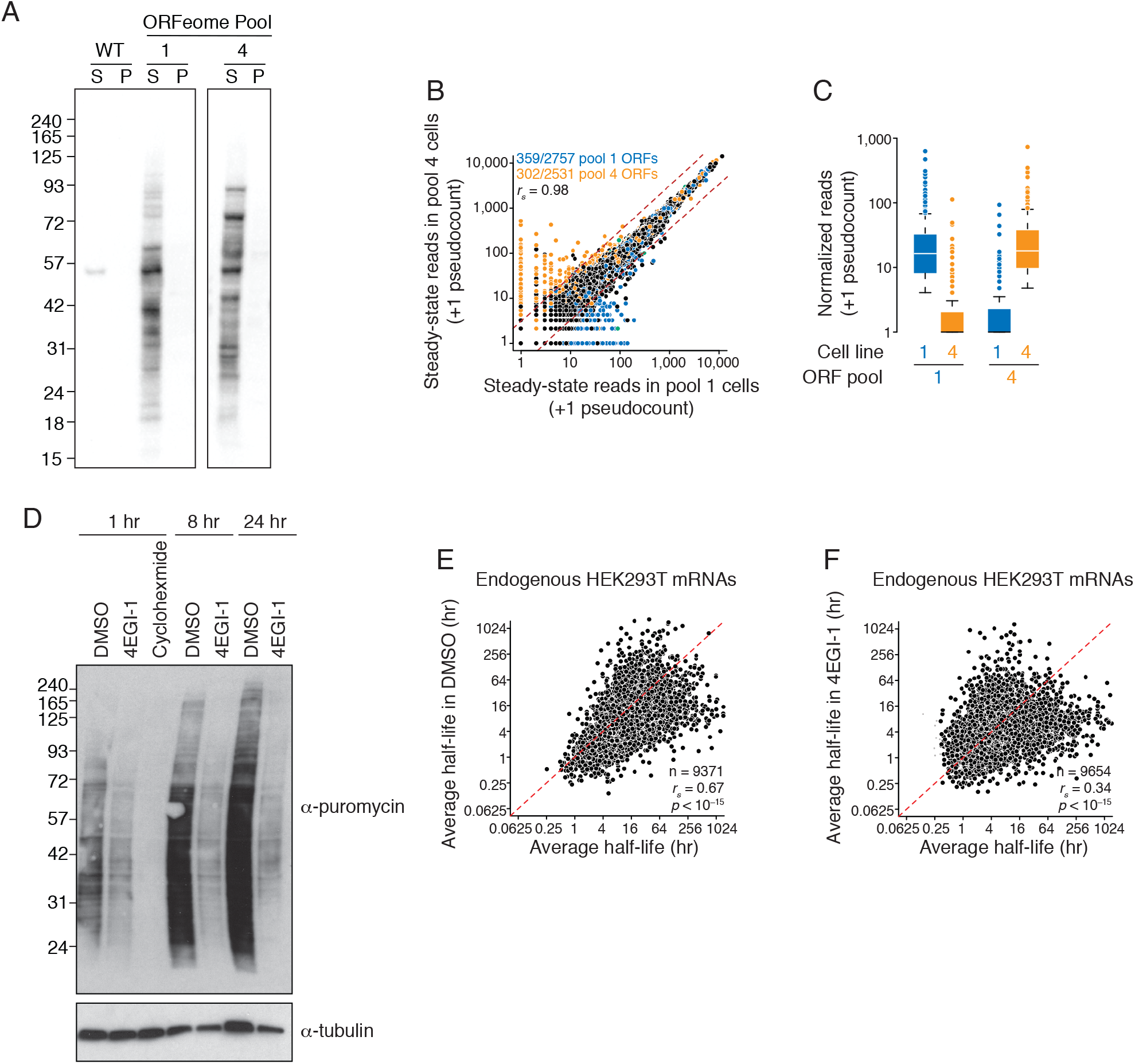
Validation of the ORFeome lines, related to Figure 2. (A) ORFeome complexity was maintained through stable cell line generation. Shown is a western blot probing lysates from the pooled ORFeome stable lines with V5 (the common C-terminal tag in the ORFeome collection). WT, parental HEK293T line; S, supernatant; P, pellet. (B) ORFeome-derived mRNAs are expressed in the stable cell lines. Shown is a scatter plot comparing steady-state RNA-seq reads (with a +1 pseudocount) for each gene between the two pooled lines used in this study. In black, genes in neither pool; in blue, genes in pool 1; in orange, genes in pool 4; in green, genes in both pools. Red dashed lines represents y = 3X and y = X/3, which were used as cut-offs to classify genes as ORFeome-expressed. ORFeome genes that did not pass threshold were not used for subsequent analysis (see Methods for more details). Numbers refer to the total number of genes in each pool and the number passing the 3-fold threshold. (C) ORFeome mRNAs are expressed in a pool-dependent fashion. Shown are boxplots of normalized read counts (with a +1 pseudocount) for ORFeome-derived mRNAs (split into pool 1 and pool 4) in the two pooled stable cell lines. Abundance in cell line 1 is shown in blue; in cell line 4, in orange. Note that the ORFeome pools are expressed in the appropriate cell line. (D) 4EGI-1 substantially reduces translation. Cells were treated with DMSO, 4EGI-1, or cyclohexamide (as a positive control) for the indicated times and then briefly treated with puromycin, which is incorporated into nascent peptides. Lysates were separated by gel electrophoresiss and probed for puromycin (top) or tubulin (bottom). (E) DMSO treatment does not substantially affect mRNA stability. Shown are scatterplots comparing half-lives for endogenous genes (averaged from both pools) from the original experiment and DMSO-treated cells. Red dashed line represents x = y. (F) 4EGI-1 treatment affects mRNA stability. As in E, except comparing half-lives from the original experiment and 4EGI-1-treated cells.

**Figure S3.**
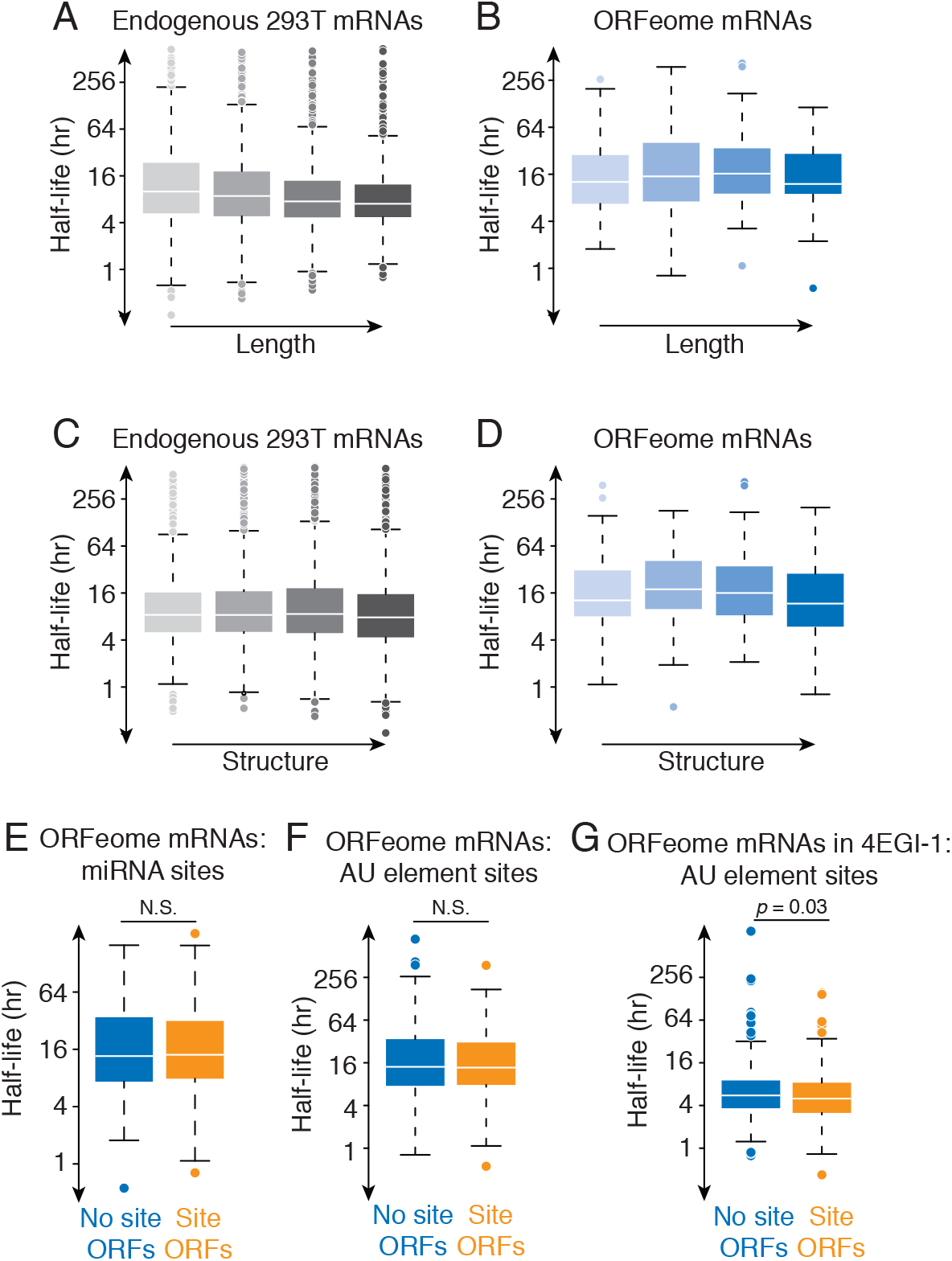
Variation in ORFeome mRNA stabilities cannot be explained by length, local secondary structure, and RBP-binding sites, related to Figure 3. (A) Endogenous mRNA stability negatively correlates with length. Shown are boxplots for half-lives of endogenous HEK293T mRNAs binned into quartiles by ORF length. (B) ORFeome mRNA stability does not correlate with length. As in A, except for ORFeome mRNAs. (C) Endogenous mRNA stability weakly correlates with local secondary structure. For each ORF, the folding energy in 100 bp sliding windows was calculated, and the minimum value taken. Shown are boxplots for half-lives of endogenous HEK293T binned into quartiles by folding energy (with increased secondary structure on the right). (D) ORFeome mRNA stability does not correlate with local secondary structure. As in B, except for ORFeome mRNAs. (E) microRNA-mediated regulation cannot explain the variation in ORFeome stability. ORFs were classified as containing or lacking seed-matched sites for the top five expressed mRNAs (site ORFs [orange] and no site ORFs [blue], respectively). Shown are boxplots for their half-lives. Significance was calculated by the Kolmogorov-Smirnov test. (F) AU-rich elements cannot explain the variation in ORFeome stability. As in E, except for AU-rich elements. (G) AU-rich elements in ORFs destabilize mRNAs upon translational repression. As in F, except for half-lives determined in the presence of 4EGI-1.

**Figure S4.**
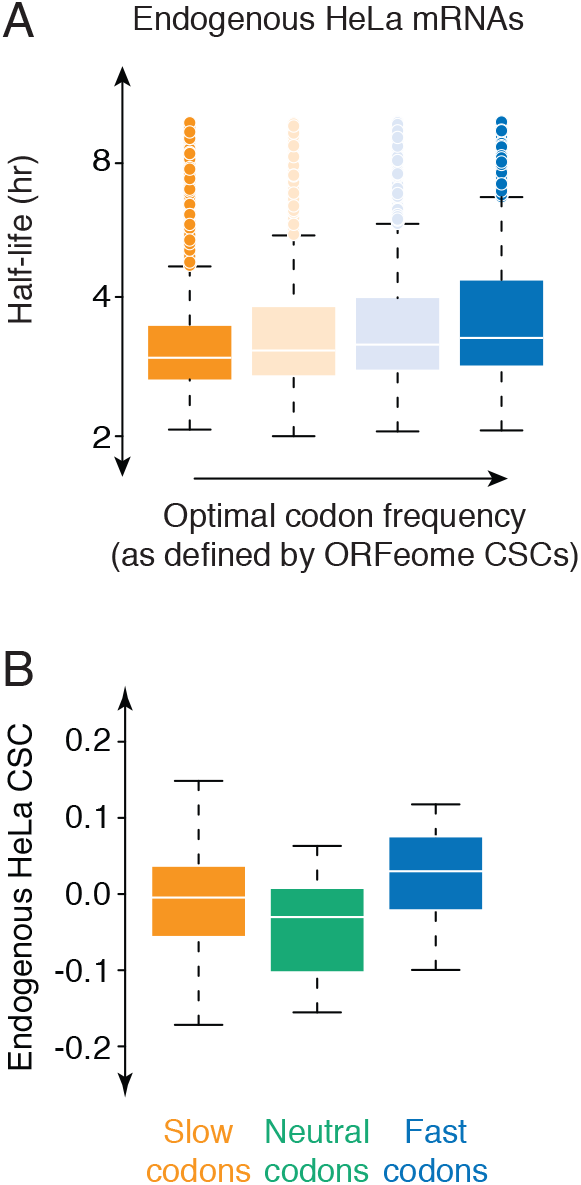
The relationship between HeLa mRNA codon content, mRNA stability, and translation elongation, related to Figure 3. (A) HeLa mRNAs with more optimal codons are more stable. Shown are boxplots of mRNA half-lives for HeLa mRNAs, binned into quartiles by the frequency of optimal codons (as determined by ORFeome-derived mRNAs). The line represents the median half-life, and the box, 1^st^ and 3^rd^ quartiles. (F) Endogenous HeLa CSCs do not correspond with pause scores. Using HeLa ribosome profiling, pause scores were calculated for each codon in the A site, and then codons were divided into three groups (slow in orange; neutral in green; fast in blue). Shown are boxplots for the corresponding CSC values as determined by endogenous HeLa mRNAs.

**Figure S5.**
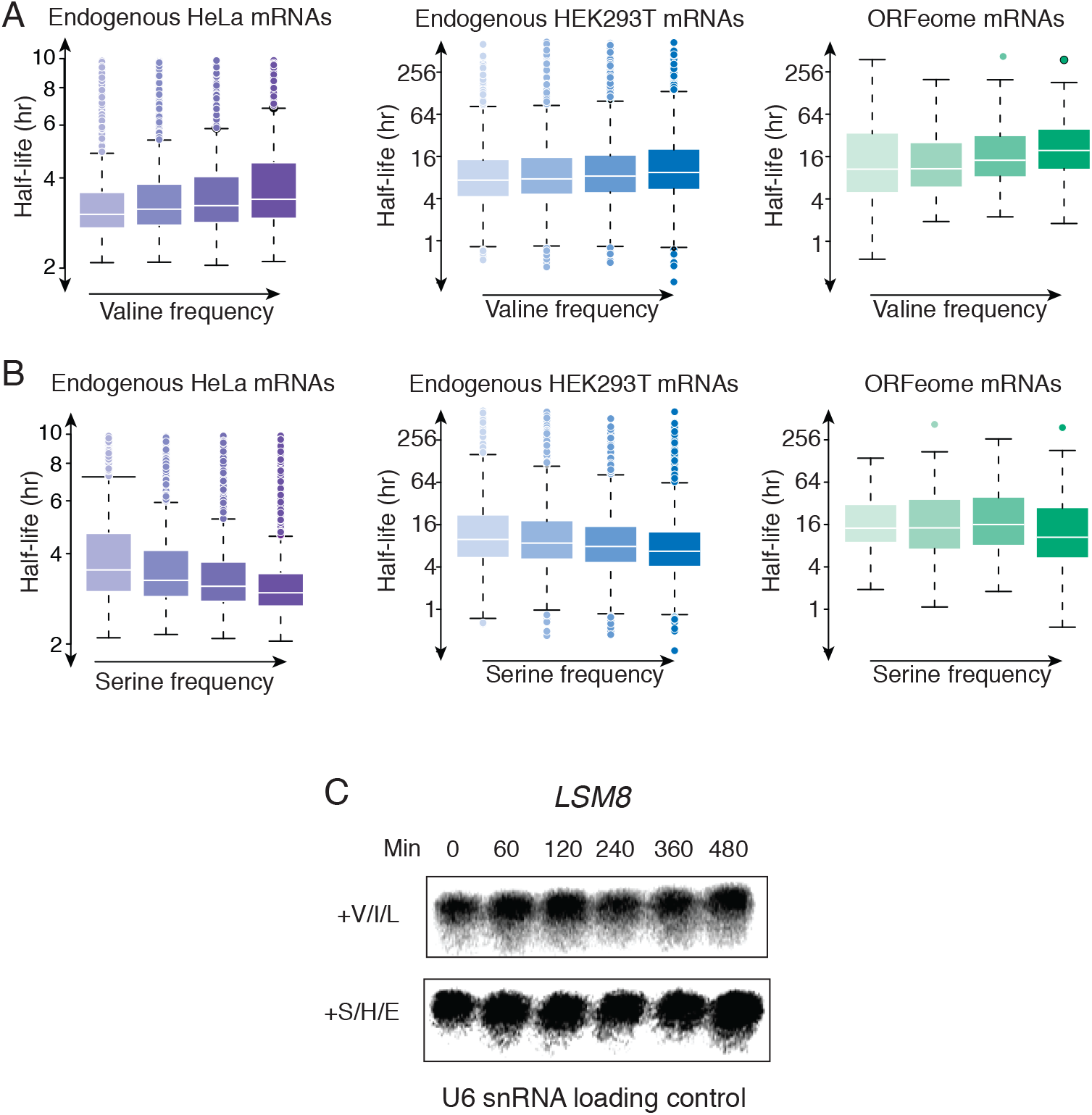
Amino acid frequency impacts mRNA stability, related to Figure 4. (A) Valine frequency correlates with mRNA stability. Shown are boxplots of mRNA stabilities for HeLa, endogenous HEK293T, and ORFeome mRNAs binned into quartiles by valine frequency. (B) Serine frequency negatively correlates with mRNA stability. As in A, except for serine. (C) U6 snRNA northern analysis for transcription shut-off experiments for the *LSM8* variants shown in Figure 1C. (D) U6 snRNA Northern analysis for *LSM8* variable amino acid content transcription shutoff/Northern mRNA decay analyses shown in Figure 4I.

**Figure S6.**
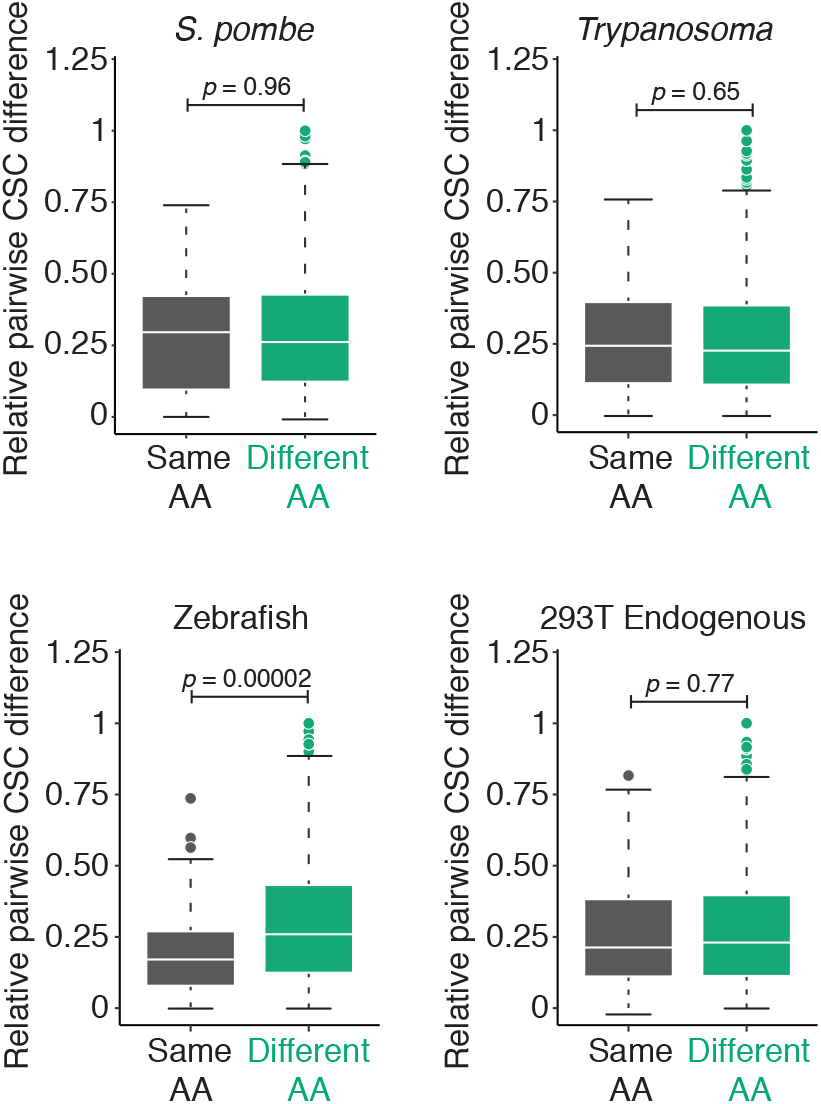
The relative influence of codon and amino acid usage varies between different organisms, related to Figure 6. For each pair of codons, the absolute difference in the corresponding CSC values was calculated and then normalized to the maximal difference (to correct for differences in overall variance between organisms). Pairs of codons were binned into these encoding the same or different amino acid (n = 87, in grey, and n = 1742, in green, respectively). Shown are boxplots of those differences for *S. pombe*, trypanosomes, zebrafish, and endogenous HEK293T mRNAs. Significance determined by Wilcoxon rank sum test.

**Table S1. Oligonucleotide and gBlock sequences used in this study, related to Figures 1 and 4.**

**Table S2. Plasmids used in this study, related to Figures 1, 2, and 4.**

**Table S3. Half-life datasets and FASTA coding sequences for additional organism CSC calculations, related to Figure 6 and Figure S6.**

## STAR METHODS

### CONTACT FOR REAGENT AND RESOURCE SHARING

Please direct any requests for further information or reagents to the lead contact, Olivia Rissland.

## EXPERIMENTAL MODEL AND SUBJECT DETAILS

### Cell lines and growth conditions

#### Human cell lines

HEK293 Tet-Off^®^ cells (Clontech 631152) were maintained in DMEM, high glucose, pyruvate (Thermo Fisher Scientific Cat #11995065) supplemented with 10% FBS (Gibco 1992275) and 1% Penicillin/Streptomycin. Human HEK293T cells were cultured in DMEM (Lonza) supplemented with 10% fetal bovine serum (FBS) (VWR Seradigm) and 1% penicillin-streptomycin solution (BioShop). Cell lines were cultured at 37°C in a humidified incubator with 5% CO2. All cells were cultured in Greiner Cellstar^®^ cell culture dishes.

#### Drosophila cell lines

*Drosophila melanogaster* Schneider 2 (S2) cells (Thermo Fisher Scientific R69007) were cultured in ExpressFive SFM media (Thermo Fisher Scientific) supplemented with 10% heat-inactivated FBS (Wisent) and 20mM L-Glutamine (Life Technologies) at 28°C.

#### Yeast strains

*S. cerevisiae* USY006 was grown in YPD liquid or plates at 30°C. Cultures were obtained from Dr. John Rubinstein (The Hospital for Sick Children Research Institute).

## METHOD DETAILS

### Plasmids and oligonucleotides

All oligonucleotides used in this study are listed in Table S1. All plasmids used in this study are listed in Table S2.

### ORFeome cell line preparation

The human ORFeome collection version 8.1 (ccsbBroad304) cloned into lentiviral vector pLX304 was obtained from Dr. Jason Moffat (University of Toronto) as a series of 96-well overnight bacterial cultures. Equal volumes of bacterial cultures were pooled into 36 pools comprising ~576 clones each. Plasmid DNA was isolated using the GeneJET Plasmid Midiprep Kit (Thermo Scientific) as per manufacturer’s instructions, yielding an average of ~70 μg plasmid DNA per pool. The 36 isolated pools were further combined into 6 unique pools for downstream cell line generation.

Each of the 6 unique virus pools were packaged by transfection into HEK293T cells using lipofectamine 2000 (Life Technologies), according to manufacturer’s instruction. Cells were transfected with the lentivirus pLX304 pool, psPAX2 packaging vector, and pVSV-G envelope vector. After 8 hours, transfection media was then removed and switched to harvest media (DMEM + 10% FBS + 1.1 g/100ml BSA (7.5% solution, Life Technologies)). Cells were left for 2 days to complete virus production.

Media was then collected from the plate and filtered through a 0.45 μm filter (Acrodisc) by syringe. Harvested viruses were aliquoted.

Freshly thawed HEK293T cells were grown in 10 cm dishes to reach ~30-50% confluence for the day of infection. Media was removed and 9 ml pre-warmed infection media (DMEM + 10% FBS + 8 μg/mL Polybrene) was added to cells. 2 ml of freshly harvested virus pool were added to 1 plate of HEK293T cells each and incubated overnight. Cells were then trypsinized and expanded into 15cm dishes. Cells were selected using selection media (DMEM + 10% FBS + 6 μg/mL Blasticidin (BioShop)) for 6 days. Selection media was changed every day. Cells were then frozen in cell freezing medium (Sigma-Aldrich) and stored in liquid nitrogen.

### Western blotting

Cells were harvested by trypsinization and pelleted by centrifugation at 1,000G for 2 minutes at 4°C. Cell pellets were resuspended in 500 μl lysis buffer (100 mM KCl, 0.1 mM EDTA, 20 mM HEPES-KOH pH 7.6, 0.4% NP-40, 10% glycerol, 1 mM DTT, complete mini EDTA-free protease inhibitors (Roche)) and clarified at 21,000G for 5 minutes at 4°C. 250 μl supernatant was mixed with 20μL 4x Bolt LDS sample buffer (Invitrogen), 8 μl 10X Bolt sample reducing agent (Invitrogen) and proteins were denatured at 75°C for 10 minutes. Protein samples were loaded into Bolt 4-12% Bis-TRIS Plus gels (Invitrogen) and run at 160V for ~1 hour. The gel was transferred onto an Amersham Hybond PVDF membrane (GE Healthcare) at 20V for ~1 hour and blocked in PBST (1X PBS with 1% Tween-20 (Sigma-Aldrich)) with 5% milk (BioBasic) for 30 minutes. Primary antibodies were added at 1: 10,000 concentration for α-V5 antibodies (Sigma-Aldrich V8012), 1: 10,000 for α-puromycin antibodies (Kerafast 3RH11), and 1: 5,000 for α-tubulin antibodies (Sigma-Aldrich T5168). Blots were incubated shaking in primary antibody overnight at 4°C.

Blots were then washed 3x with PBST for 5 minutes each and incubated with 1: 10,000 concentration α-mouse secondary antibody (NEB 7076) for 1 hour at room temperature. Blots were washed with PBST 3X for 5 minutes each. Blots were imaged using ECL Prime Western Blotting Detection Reagent (GE Healthcare) and exposed on Amersham Hyperfilm (GE Healthcare).

### Polysome fractionation

hORF cell line 1 was grown for 24 hours in the presence of either DMSO or 100 μM 4EGI-1 (Cedarlane). Cells were treated with 100 μg/ml cycloheximide (CHX) (BioShop) for 10 minutes. Cells were harvested on ice by washing 2x with ice-cold PBS containing 100 μg/ml CHX, and lysing with 500 μl ice-cold filter-sterilized lysis buffer (10 mM Tris-HCl (pH 7.4), 5 mM MgCl_2_, 100 mM KCl, 1% Triton X-100, 2 mM DTT, 500 U/ml RNasin (Promega), 100 μg/ml CHX, Protease inhibitor (1X complete, EDTA-free, Roche)). Cells were scraped off the dish into tubes and sheared gently 4x with a 26-guage needle. Lysed cells were centrifuged at 1,300G for 10 minutes at 4°C, and clarified supernatant was isolated.

A 10/50% sucrose gradient was created by combining heavy and light solutions on a BioComp Gradient MasterTM. Heavy and light solutions consisted of 20 mM HEPES-KOH (pH 7.4), 5 mM MgCl_2_, 100 mM KCl, 2 mM DTT, 100 μg/ml CHX, and 20 U/ml SUPERaseIn, and either 10% or 50% sucrose (w/v) respectively. 300 μl of samples were layered on sucrose gradients and centrifuged in a pre-cooled Beckman Ultracentrifuge L-90K using SW41 rotor at 36,000 RPM (221632.5G) for 2 hours at 4°C. The gradient was fractionated using the BioComp Piston Gradient FractionatorTM and absorbance measurements were made using an Econo EM-1 UV Monitor (BioRad).

### Puromycin incorporation assay

hORF cell line 1 was grown for 1, 8, or 24 hours in the presence of either DMSO, 100 μM 4EGI-1, or 5 μg/ml cycloheximide. Cells were pulsed with 1.5 μg/ml puromycin dihydrochloride (Gibco) for 10 minutes at 37°C. Cells were then harvested and lysed as above and probed by western blot with α-puromycin antibody (Kerafast 3RH11) to detect overall incorporation.

### HEK293T endogenous and ORFeome mRNA stability determination by metabolic labeling

#### Generation of spike-in RNA

Two sets of spike-in RNA were generated. An unlabeled *S. cereivisae* spike-in is used to determine the enrichment of 4SU-labeled RNA over unlabeled RNA, as described previously (Lugowski et al., 2017). *S. cereivisae* strain USY006 was grown in YPD liquid culture at 30°C, and RNA was isolated using hot acidic phenol method (Rissland and Norbury, 2009). A 4SU-labeled *D. melanogaster* spike-in was also generated by supplementing S2 culture media with 100 μM 4SU for 24 hours prior to harvesting. RNA was extracted using TRI-reagent (Molecular Research Center) as per manufacturer’s instructions.

#### Metabolic labeling of hORFeome cell lines

Freshly thawed HEK293T hORFeome cell lines were cultured for 3-4 passages and seeded into DMEM + FBS culture media in 15 cm dishes such that they attained ~50% confluence on the day of the time course. Media was replaced with DMEM + 10% FBS + 100 μM 4SU (Sigma-Aldrich) reconstituted in DMSO. Cells were harvested at 1, 2, 4, 8, 12, and 24 hours after addition of 4SU. Harvesting was performed by dislodging cells off the plate during two washes with cold 1X PBS followed by spinning at 1,000G for 5 minutes at 4°C. Cell pellets were resuspended in 1mL TRI-Reagent (Molecular Research Center) and extracted according to manufacturer instructions.

For translation inhibition experiments, hORF cell line 1 or cell line 4 cells growing in 10mL DMEM + FBS in 10cm dishes were pre-treated with either 0.1% DMSO or 100μM 4EGI-1 (Cedarlane) dissolved in DMSO for 1 hour. Following this, 100μM 4SU was added to media for all plates, and the time course was performed as described above.

#### Reversible biotinylation and fractionation of 4SU-labeled mRNAs

RNA was labeled as described previously (Lugowski et al., 2017). Briefly, 100 μg of total hORF RNA was mixed with 10 μg unlabeled *S. cerevisiae* RNA (i.e., 10% w/w) and 10 μg 4SU labeled S2 *D. melanogaster* RNA (i.e., 10% w/w). Water was added to bring the volume up to 120 μl. 1 mg/mL HPDP-biotin (Thermo Fisher Scientific) was reconstituted in dimethylformamide by shaking at 37°C for 30 minutes at 300 RPM. 120 μl of 2.5x Citrate buffer (25 mM citrate, pH 4.5, 2.5 mL EDTA) and 60 μl of 1 mg/mL HPDP-biotin were added to the pre-mixed RNA sample for each time point. This solution was incubated at 37°C for 2 hours at 300 RPM on an Eppendorf ThermoMixer F1.5 in the dark. Samples were extracted twice with acid phenol, pH 4.5 (Invitrogen), and once with chloroform. RNA was precipitated with 18 μl 5M NaCl, 750 μl 100% ethanol, and 2 μl GlycoBlue (Invitrogen) overnight at -20°C. Precipitated RNA was pelleted for 30 minutes at 21,000G at 4°C. The RNA pellet was resuspended in 200 μl of 1x wash buffer (10 mM Tris-Cl, pH 7.4, 50 mM NaCl, 1 mM EDTA).

Biotinylated RNA was purified using the μMACS Streptavidin microbeads system (Miltenyi Biotec). 50 μ; Miltenyi beads per sample were pre-blocked with 48 μl 1x wash buffer and 2 μl yeast tRNA (Invitrogen), rotating for 20 minutes at room temperature. μMACS microcolumns were washed 1x with 100 μl nucleic acid equilibration buffer (Miltenyi Biotec), followed by 5x washes with 100 μl 1x wash buffer. Beads were applied to microcolumns in 100μL aliquots, and again washed 5x with 100 μ; 1x wash buffer. Beads were demagnetized and eluted off the column with 2x 100 μl 1x wash buffer, and columns were placed back on the magnetic stand. 200 μl beads were mixed with each sample of biotinylated RNA and rotated at room temperature for 20 minutes.

Samples were then applied to the microcolumns in 100 μl aliquots, washed 3x with 400 μl wash A buffer (10mM Tris-Cl, pH 7.4, 6M urea, 10mM EDTA) pre-warmed to 65°C, and then washed 3x with 400 μl wash B buffer (10mM Tris-Cl, pH7.4, 1M NaCl, 10mM EDTA). RNA was eluted with 5x 100 μl of 1x wash buffer supplemented with 0.1M DTT, and flow through was collected in a tube. Purified RNA was precipitated with 30 μl 5M NaCl, 2 μl GlycoBlue, and 1ml 100% ethanol, incubated at –20°C overnight. Samples were then spun at 21,000G at 4°C for 30 minutes and resuspended in 20 μl water. RNA quality was assessed by running 3 μl of samples on a ~1.5% agarose gel.

#### Generation of next generation sequencing libraries and RNA-sequencing

10 μl of purified 4SU-labeled RNA or unpurified total RNA from the 24-hour time point was used to prepare RNA-seq libraries using the TruSeq Stranded mRNA Sample Preparation Kit (Illumina), according to manufacturer’s instructions. Adapter-ligated fragments were enriched with 14x PCR cycles. ~16-22 samples were multiplexed on a single lane in an Illumina HiSeq 2500 at The Centre for Applied Genomics (TCAG, SickKids) to obtain ~10 million 50bp single-end reads per sample.

### Single gene mRNA decay analysis

Plasmid DNA was transfected into HEK293 Tet-Off^®^ cells using Lipofectamine 2000 (Thermo Fisher Scientific 11668027), according to manufacturer’s instructions. Transfection media was removed after 24 hours. 48 hours post-transfection, media was replaced with DMEM + Tet-free FBS + 2 ug/ml doxycycline (Sigma-Aldrich Cat# D3072). Cells were harvested using 1 ml Trizol (Thermo Fisher Scientific 15596018) at indicated time points, and RNA isolated according to manufacturer’s instructions. For steady-state mRNA analyses, cells were plated and transfected as above; cells were harvested using 1 mL Trizol (Thermo Fisher Scientific 15596018) 48 hours post-transfection.

Agarose northern analyses were performed as described in (Presnyak et al., 2015). Briefly, RNA was loaded onto 1.4% formaldehyde agarose gels and run at 100V for 90 min. Gels were imaged to check for ribosomal RNA quality and quantity before blotting onto Hybond-N+ membrane (GE Amersham RPN303B) by overnight transfer by capillary action.

All Firefly luciferase, *MECP2*, *CFTR*, and *LSM8* reporter transcripts were detected using a ^32^P-α-CTP (Perkin-Elmer; 3000 Ci/mmol) radiolabeled asymmetric PCR probe (oJC3609/10; see Table S1) directed towards the pJC842 synthetic 3’UTR. Probe was hybridized overnight at 65°C, then washed twice using 2X SSC/0.1% SDS at 24°C (5 min each) followed by washing in 0.1% SSC/0.1% SDS at 50°C for 1 hr.

U6 snRNA Northern probe was created by end-labeling U6 snRNA oligo probe (see Table S1) with γ-^32^P-ATP (Perkin Elmer; specific activity = 6000 Ci/mmol) using T4 Polynucleotide Kinase (New England Biolabs, Cat. M0201S). The probe was hybridized overnight at 42°C, then washed twice in 6X SSC/0.1% SDS at 24°C for 15 min, then at 50°C for 15 min.

Blots were exposed on a storage phosphor screen for 15 min (U6 snRNA) or overnight (Firefly, *MECP2, CFTR*, and *LSM8* reporters). Stored signal was read using the Typhoon 9400 Variable Mode Imager (Amersham Biosciences). Quantitation of phosphorimager signal was performed using ImageQuant software (Molecular Dynamics; version 5.2).

Polyacrylamide Northern analysis of *LSM8* reporter mRNA was performed using 6% polyacrylamide/urea denaturing gels. Samples were run at 400V for 16 hr in 1X TBE and transferred at 50V in 0.5X TBE for 3 hours at 4°C. Hybridization with radiolabeled asymmetric PCR probe, washing, and detection proceeded as described above.

## QUANTIFICATION AND STATISTICAL ANALYSIS

### HeLa mRNA half-life data analysis

Processed BRIC-Seq half-life data for HeLa cells was obtained from the Gene Expression Omnibus (Accession: GSE102113; see (Arango et al., 2018) for alignment, filtering, and half-life calculation details). A tidy dataframe was constructed using transcripts with reported half-lives for wild-type HeLa cells. Human coding sequences from genome build GRCh38 were obtained from Ensembl (ftp://ftp.ensembl.org/pub/release-94/fasta/homo_sapiens/dna/); ORFs were restricted to annotated coding sequences starting with “ATG”. CSC/AASC calculations were performed using wild type HeLa half-lives as described below.

### HEK293T endogenous and ORFeome mRNA half-life Calculations

*Reference genome information*. Human (hg38), *D. melanogaster* (dm6), and *S. cerevisiae* (sacCer3) genomes were obtained in 2bit format using the UCSC Table Browser (Karolchik et al., 2004). 2bit files were converted to FASTA using the kentUtils command twoBitToFa, and GTF annotations were downloaded using the kentUtils command genePredToGtf. The three genomes were combined using custom bash scripts to make a hg38+dm6+sacCer3 genome.

### Initial processing of sequencing reads

Library quality was assessed using FastQC v0.11.5 (http://www.bioinformatics.babraham.ac.uk/projects/fastqc). Reads were trimmed and clipped for Illumina adapters using Trimmomatic v0.36 (Bolger et al., 2014) using the following settings: -phred33 ILLUMINACLIP: TruSeq3-SE.fa:2:30:10 LEADING:3 TRAILING:3 SLIDINGWINDOW:4:15 MINLEN:36.

#### Genome mapping and counting

Trimmed reads were aligned to the indexed hg38+dm6+sacCer3 genome using STAR version 2.5.2 (Dobin et al., 2013) with the following non-default settings: --outFilterMultimapNmax 10 -- outFilterMismatchNoverLmax 0.05 --outFilterScoreMinOverLread 0.75 -- outFilterMatchNminOverLread 0.85 --alignIntronMax 1 --outFilterIntronMotifs RemoveNoncanonical --outSAMtype BAM SortedByCoordinate --quantMode GeneCounts.

HTSeq version 0.6.1 (Anders et al., 2015) was then used to quantify gene counts from aligned BAM files using the following settings: --order=pos --stranded=reverse --minaqual=10 --mode=intersection-strict. Note that counting of features from human CDS and intronic GTF files were performed separately.

#### Defining hORF genes

Gene counts were loaded into RStudio version 3.3.1. Dplyr package (https://CRAN.R-project.org/package=dplyr) was used for all data manipulation and filtering. Steady state RNA-sequencing counts mapping to human CDS features were obtained from each cell line and normalized to library size to allow comparisons across samples. For a given cell line X, genes were described as “detectable hORF genes” if they met each of the following conditions:

1. They were in the list of hORFs infected into cell line X;
2. Normalized steady state RNA-sequencing for cell line X was greater than 3-fold that in the other cell line;
3. Normalized steady state RNA-sequencing for cell line X was greater than 4 reads

Genes that were not infected into cell line X were described as endogenous genes.

#### Calculation of mRNA half-lives

All half-life calculations were performed in RStudio version 3.3.1 as described previously (Lugowski et al., 2018). Briefly, read counts for mature human mRNAs were filtered such that each gene had at least 1 read mapped to its CDS at each time point, and at least 5 reads mapped at any (at least 1 of 6) time point. CDS-mapping reads for each gene at each time point were then normalized to the sum of all corresponding *D. melanogaster* mapping reads.

Half-lives were calculated by fitting these normalized read counts at each time point to a bounded growth equation using weighted nonlinear least squares. The bounded growth equation has been previously described (Lugowski et al., 2017). Briefly, the equation states:

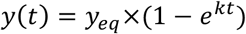

where y(t) is the amount of a given transcript remaining at time t, y_eq_ is the amount of that transcript at steady state, and k is the transcript-specific decay constant.

The nls() function in the stats package was used to fit the time points to the equation above, with settings equivalent to the following:

- start = c(y_eq_ = max(y), k = -0.5)
- algorithm = “port”
- weights = 1/y(t)
- lower = c(y_eq_ = 0, k = -Inf), upper = c(y_eq_ = Inf, k = 0)

If the data did not converge, a value of NA was returned. The half-life of each transcript is then obtained using the following equation:

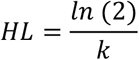

### Calculation of local secondary structure

To measure local secondary structure, each gene’s coding sequence was assayed by sliding 100bp windows, each starting 3bp apart. Each window was folded using ViennaRNA version 2.2.8 RNAfold function (Lorenz et al., 2011) using default parameters. Minimum folding energy (MFE) for each 100bp sequence was extracted from output files using custom bash scripts. Median and minimum MFEs across each CDS were determined using group_by and summarize functions in the dplyr package in RStudio.

### Codon and amino acid stability coefficient calculations

Codon usage frequency was calculated from each gene’s coding sequence using the seqinr package’s uco function. Amino acid usage was calculated using the translate function in the Biostrings R package to convert coding sequence into amino acid sequence, and then using Biostrings alphabetFrequency function to count amino acid per CDS. Amino acid usage frequency was determined by dividing amino acid count by CDS length. For frame shift controls, codon and amino acid usage were calculated after shifting the frame by +1 (removing positions 1, n-2, and n-1 from CDS of length n) and +2 (removing positions 1, 2, and n-1 from CDS of length n).

As described previously (Radhakrishnan et al., 2016), codon stability coefficients (CSC) from a given half-life dataset were calculated by determining the Spearman correlation between the codon frequency for each codon in a transcript with the measured half-lives of that transcript. Stop codons were excluded from CSC calculations. AASCs were similarly calculated by determining the Spearman correlation between the frequencies for each amino acid in a transcript with the measured half-lives of that transcript.

### Codon and amino acid-specific pause score calculations using HeLa Ribo-Seq

Processed Ribo-seq BAM files for HeLa cells was obtained from the Gene Expression Omnibus (Accession: GSE102113; see Arango et al. 2018 for details of alignment and filtering). As described in Arango et al. 2018, A-site codons were predicted based on position and total length of each ribosome-protected fragment read relative to the annotated start and stop codons of transcripts (under the assumption that reads were in frame within their respective ORF sequences) using an offset of 17 nt from the 5’ end of the ribosome-protected fragment read. Codon-specific ribosome pause scores were expressed as average ribosome density per codon (Ribo-seq reads per codon / total Ribo-seq reads over the entire annotated ORF density) using a bootstrap approach, where the average ribosome density represents the 50^th^ percentile of 5000 iterations. Codon-specific ribosome pause scores with respect to the P-site and E-site codon were determined in an analogous manner, except the centering frame offset was shifted to 14 (P site) or 11 (E site) in place of default 17 (A site). Amino acid-specific ribosome pause scores were determined by taking the arithmetic mean of codon-specific pause scores for groups of synonymous codons (n=1 to n=6).

### CSC difference calculations for additional organisms

CSC calculations were performed as described above using half-life datasets and FASTA-formatted coding sequences indicated in Table S3. Following CSC calculations, all possible codon pairings were generated using the Python 3 itertools package; CSC values were matched to each codon. Relative CSC differences were calculated for each codon pair as follows:

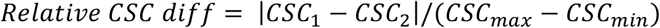

where *CSC_max_* = maximum observed CSC and *CSC_min_* = minimum observed CSC for any given organism.

### Gene group AASC analysis

Gene symbols for various protein families or related pathways were retrieved from the Gene Ontology Consortium (htpp://geneontology.org/; TCA cycle: GO:0006099; ATP synthesis: GO:0006754; Glycolysis: GO:0006096; Splicing proteins: GO:0000398; Cell cycle proteins: GO:0007049), the InterPro Protein Sequence Analysis and Classification database (http://www.ebi.ac.uk/interpro/; Protein kinase family: IPR017892; KH Domain protein family: IPR004088), and the HUGO Gene Nomenclature Committee (http://genenames.org; Zinc finger proteins Group ID: 26; Serine/Arginine-rich proteins Group ID: 737; Cytoplasmic ribosomal proteins Group IDs: 728 + 729; Mitochondrial ribosomal proteins Group ID: 646).

For the purposes of gene groups analysis, additional cytoplasmic ribosomal protein transcripts (HGNC Group IDs 728 + 729) were identified which otherwise passed filtering (i.e. adequate read depth across the entire time course for both HeLa BRIC-Seq replicates), yet failed half-life estimation by linear modeling due to high stability; these transcripts were manually assigned the maximum half-life (24 hrs) and added to existing half-life data. Weighted average transcript AASC was calculated as follows:

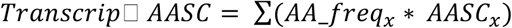

where *x* = any given amino acid.

For cytoplasmic and mitochondrial ribosomal proteins, weighted CSC scores for all synonymous codons were calculated in an analogous manner:

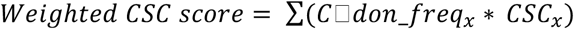

where x = a given synonymous codon.

### Other statistical analyses

Number of replicates, statistical tests used, and *p*-values are specified in the figures and figure legends.

## DATA AND SOFTWARE AVAILABILITY

The accession number for the raw data files for the reported in this paper is NCBI Gene Expression Omnibus GSE123165. The accession number for the raw and processed HeLa BRIC-Seq data and HeLa ribosome sequencing data used in this paper is NCBI Gene Expression Omnibus GSE102113. All half-life datasets and coding sequence files used for additional organisms in this study are listed in Table S3.

---

KEY RESOURCES TABLE.

KEY RESOURCES TABLE
| REAGENT or RESOURCE | SOURCE | IDENTIFIER |
| --- | --- | --- |
| **Antibodies** |  |  |
| Anti-Puromycin | Kerafast | 3RH11 |
| Anti-V5 | Sigma-Aldrich | V8012 |
| Anti-Tubulin | Sigma-Aldrich | T5168 |
| Anti-mouse IgG, HRP-linked | New England Biolabs | 7076 |
| **Bacterial and Virus Strains** |  |  |
| N/A |  |  |
| **Biological Samples** |  |  |
| N/A |  |  |
| **Chemicals, Peptides, and Recombinant Proteins** |  |  |
| 4EGI-1 | Cedarlane | 4800 |
| 4-Thiouridine (4SU) | Sigma-Aldrich | T4509 |
| Cycloheximide | BioShop Canada | CYC003 |
| **Critical Commercial Assays** |  |  |
| TruSeq mRNA Stranded mRNA Sample Preparation Kit | Illumina | 20020594 |
| **Deposited Data** |  |  |
| Human reference genome NCBI build 38, GRCh38 | Genome Reference Consortium | http://hgdownload.soe.ucsc.edu/goldenPath/hg38/bigZips/hg38.2bit |
| *Saccharomyces cerevisiae* reference genome build R64, sacCer3 | Saccharomyces Genome Database | http://hgdownload.soe.ucsc.edu/goldenPath/sacCer3/bigZips/sacCer3.2bit |
| *Drosophila melanogaster* reference genome release 6.18, dm6 | The FlyBase Consortium/Berkeley Drosophila Genome Project | http://hgdownload.soe.ucsc.edu/goldenPath/dm6/bigZips/dm6.2bit |
| HEK293T endogenous raw and processed half-life data | This paper | GEO: GSE123165 |
| ORFeome collection raw and processed half-life data | This paper | GEO: GSE123165 |
| HeLa BRIC-Seq processed half-life data, wild type | Arango et al. 2018 | GEO: GSE102113 |
| HeLa processed ribosome sequencing data | Arango et al. 2018 | GEO: GSE102113 |
| Table S3: Additional half-life data and FASTA formatted coding sequences for additional organisms (*S. cerevisiae*, *S. pombe*, *D. melanogaster*, *D. rerio*, *T. brucei*) | Presnyak et al. 2015 | GEO: GSE57385 |
| **Experimental Models: Cell Lines** |  |  |
| HEK293 Tet-off(R) Advanced Cell Line | Clontech | 631152 |
| HEK293T | ATCC | CRL-3216 |
| Human ORFeome cell lines, pools 1 and 4 | This paper | N/A |
| **Experimental Models: Organisms/Strains** |  |  |
| Yeast USY006 | John Rubenstein | N/A |
| **Oligonucleotides** |  |  |
| Table S1: gBlocks and Northern probe sequences | Integrated DNA Technologies (IDT) | N/A |
| **Recombinant DNA** |  |  |
| Table S2: Firefly luciferase, *MECP2*, *CFTR*, and *LSM8* Reporters; lentiviral packaging vectors | N/A | N/A |
| Human ORFeome V8.1 Library | Yang, et al. 2011 | ccsbBroad304 |
| **Software and Algorithms** |  |  |
| R | https://cran.r-project.org/bin/windows/base/old/3.4.3/ | Version 3.3.1 and 3.4.3 |
| R Biostrings | https://bioconductor.org/packages/release/bioc/html/Biostrings.html | Version 2.46.0 |
| R Tidyverse (dplyr, ggplot2, forcats, readr, stringr, purrr, tidyr, tibble) | https://cran.r-project.org/web/packages/tidyverse/index.html | Version 1.2.1 |
| FastQC | https://cran.r-project.org/web/packages/fastqcr/index.html | Version 0.11.5 |
| Trimmomatic | Bolger, et al. 2014 | Version 0.36 |
| STAR | (Dobin et al., 2013) | Version 2.5.2 |
| ImageQuant | Molecular Dynamics/ GE Healthcare | Version 5.2 |
| **Other** |  |  |
| N/A |  |  |

## REFERENCES

Anders, S., Pyl, P.T., Huber, W., 2015. HTSeq--a Python framework to work with high-throughput sequencing data. Bioinformatics 31, 166–169. doi:10.1093/bioinformatics/btu638

Arango, D., Sturgill, D., Alhusaini, N., Dillman, A.A., Sweet, T.J., Hanson, G., Hosogane, M., Sinclair, W.R., Nanan, K.K., Mandler, M.D., Fox, S.D., Zengeya, T.T., Andresson, T., Meier, J.L., Coller, J., Oberdoerffer, S., 2018. Acetylation of Cytidine in mRNA Promotes Translation Efficiency. Cell 1–40. doi:10.1016/j.cell.2018.10.030

Artieri, C.G., Fraser, H.B., 2014. Accounting for biases in riboprofiling data indicates a major role for proline in stalling translation. Genome Res. 24, 2011–2021. doi:10.1101/gr.175893.114

Bartel, D.P., 2009. MicroRNAs: target recognition and regulatory functions. Cell 136, 215–233. doi:10.1016/j.cell.2009.01.002

Bazzini, A.A., Del Viso, F., Moreno-Mateos, M.A., Johnstone, T.G., Vejnar, C.E., Qin, Y., Yao, J., Khokha, M.K., Giraldez, A.J., 2016. Codon identity regulates mRNA stability and translation efficiency during the maternal-to-zygotic transition. EMBO J. 35, 2087–2103. doi:10.15252/embj.201694699

Beelman, C.A., Stevens, A., Caponigro, G., LaGrandeur, T.E., Hatfield, L., Fortner, D.M., Parker, R., 1996. An essential component of the decapping enzyme required for normal rates of mRNA turnover. Nature 382, 642–646. doi:10.1038/382642a0

Boël, G., Letso, R., Neely, H., Price, W.N., Wong, K.-H., Su, M., Luff, J., Valecha, M., Everett, J.K., Acton, T.B., Xiao, R., Montelione, G.T., Aalberts, D.P., Hunt, J.F., 2016. Codon influence on protein expression in E. coli correlates with mRNA levels. Nature 529, 358–363. doi:10.1038/nature16509

Bolger, A.M., Lohse, M., Usadel, B., 2014. Trimmomatic: a flexible trimmer for Illumina sequence data. Bioinformatics 30, 2114–2120. doi:10.1093/bioinformatics/btu170

Burow, D.A., Martin, S., Quail, J.F., Alhusaini, N., Coller, J., Cleary, M.D., 2018. Attenuated Codon Optimality Contributes to Neural-Specific mRNA Decay in Drosophila. Cell Rep 24, 1704–1712. doi:10.1016/j.celrep.2018.07.039

de Freitas Nascimento, J., Kelly, S., Sunter, J., Carrington, M., 2018. Codon choice directs constitutive mRNA levels in trypanosomes. Elife 7, 2087. doi:10.7554/eLife.32467

Dittmar, K.A., Goodenbour, J.M., Pan, T., 2006. Tissue-Specific Differences in Human Transfer RNA Expression. PLoS Genet 2, e221. doi:10.1371/journal.pgen.0020221.st006

Dobin, A., Davis, C.A., Schlesinger, F., Drenkow, J., Zaleski, C., Jha, S., Batut, P., Chaisson, M., Gingeras, T.R., 2013. STAR: ultrafast universal RNA-seq aligner. Bioinformatics 29, 15–21. doi:10.1093/bioinformatics/bts635

Duan, J., Shi, J., Ge, X., Dölken, L., Moy, W., He, D., Shi, S., Sanders, A.R., Ross, J., Gejman, P.V., 2013. Genome-wide survey of interindividual differences of RNA stability in human lymphoblastoid cell lines. Sci Rep 3, 1318. doi:10.1038/srep01318

Eulalio, A., Huntzinger, E., Nishihara, T., Rehwinkel, J., Fauser, M., Izaurralde, E., 2009. Deadenylation is a widespread effect of miRNA regulation. RNA 15, 21–32. doi:10.1261/rna.1399509

Fabian, M.R., Cieplak, M.K., Frank, F., Morita, M., Green, J., Srikumar, T., Nagar, B., Yamamoto, T., Raught, B., Duchaine, T.F., Sonenberg, N., 2011. miRNA-mediated deadenylation is orchestrated by GW182 through two conserved motifs that interact with CCR4-NOT. Nat Struct Mol Bio 18, 1211–1217. doi:10.1038/nsmb.2149

Fabian, M.R., Frank, F., Rouya, C., Siddiqui, N., Lai, W.S., Karetnikov, A., Blackshear, P.J., Nagar, B., Sonenberg, N., 2013. Structural basis for the recruitment of the human CCR4-NOT deadenylase complex by tristetraprolin. Nat Struct Mol Bio 20, 735–739. doi:10.1038/nsmb.2572

Fabian, M.R., Mathonnet, G., Sundermeier, T., Mathys, H., Zipprich, J.T., Svitkin, Y.V., Rivas, F., Jinek, M., Wohlschlegel, J., Doudna, J.A., Chen, C.-Y.A., Shyu, A.-B., Yates, J.R., Hannon, G.J., Filipowicz, W., Duchaine, T.F., Sonenberg, N., 2009. Mammalian miRNA RISC recruits CAF1 and PABP to affect PABP-dependent deadenylation. Mol. Cell 35, 868–880. doi:10.1016/j.molcel.2009.08.004

Ferec, C., Cutting, G.R., 2012. Assessing the Disease-Liability of Mutations in CFTR. Cold Spring Harb Perspect Med 2, a009480. doi:10.1101/cshperspect.a009480

Gardin, J., Yeasmin, R., Yurovsky, A., Cai, Y., Skiena, S., Futcher, B., 2014. Measurement of average decoding rates of the 61 sense codons in vivo. Elife 3, 198. doi:10.7554/eLife.03735

Geisberg, J.V., Moqtaderi, Z., Fan, X., Ozsolak, F., Struhl, K., 2014. Global analysis of mRNA isoform half-lives reveals stabilizing and destabilizing elements in yeast. Cell 156, 812–824. doi:10.1016/j.cell.2013.12.026

Goodarzi, H., Nguyen, H.C.B., Zhang, S., Dill, B.D., Molina, H., Tavazoie, S.F., 2016. Modulated Expression of Specific tRNAs Drives Gene Expression and Cancer Progression. Cell 165, 1416–1427. doi:10.1016/j.cell.2016.05.046

Grimson, A., Farh, K.K.-H., Johnston, W.K., Garrett-Engele, P., Lim, L.P., Bartel, D.P., 2007. MicroRNA targeting specificity in mammals: determinants beyond seed pairing. Mol. Cell 27, 91–105. doi:10.1016/j.molcel.2007.06.017

Gu, S., Jin, L., Zhang, F., Sarnow, P., Kay, M.A., 2009. Biological basis for restriction of microRNA targets to the 3' untranslated region in mammalian mRNAs. Nat Struct Mol Bio 16, 144–150. doi:10.1038/nsmb.1552

Gutierrez, E., Shin, B.-S., Woolstenhulme, C.J., Kim, J.-R., Saini, P., Buskirk, A.R., Dever, T.E., 2013. eIF5A Promotes Translation of Polyproline Motifs. Mol. Cell 51, 35–45. doi:10.1016/j.molcel.2013.04.021

Hanson, G., Alhusaini, N., Morris, N., Sweet, T., Coller, J., 2018. Translation elongation and mRNA stability are coupled through the ribosomal A-site. RNA rna.066787.118. doi:10.1261/rna.066787.118

Harigaya, Y., Parker, R., 2016. Analysis of the association between codon optimality and mRNA stability in Schizosaccharomyces pombe. BMC Genomics 17, 895. doi:10.1186/s12864-016-3237-6

Herzog, V.A., Reichholf, B., Neumann, T., Rescheneder, P., Bhat, P., Burkard, T.R., Wlotzka, W., Haeseler von, A., Zuber, J., Ameres, S.L., 2017. Thiol-linked alkylation of RNA to assess expression dynamics. Nat Meth 539, 113. doi:10.1038/nmeth.4435

Ingolia, N.T., Ghaemmaghami, S., Newman, J.R.S., Weissman, J.S., 2009. Genome-wide analysis in vivo of translation with nucleotide resolution using ribosome profiling. Science 324, 218–223. doi:10.1126/science.1168978

Jeacock, L., Faria, J., Horn, D., 2018. Codon usage bias controls mRNA and protein abundance in trypanosomatids. Elife 7, 1247. doi:10.7554/eLife.32496

Karolchik, D., Hinrichs, A.S., Furey, T.S., Roskin, K.M., Sugnet, C.W., Haussler, D., Kent, W.J., 2004. The UCSC Table Browser data retrieval tool. Nucleic Acids Res. 32, D493–6. doi:10.1093/nar/gkh103

Kuzuoğlu-Öztürk, D., Bhandari, D., Huntzinger, E., Fauser, M., Helms, S., Izaurralde, E., 2016. miRISC and the CCR4-NOT complex silence mRNA targets independently of 43S ribosomal scanning. EMBO J. 35, 1186–1203. doi:10.15252/embj.201592901

Kyte, J., Doolittle, R.F., 1982. A simple method for displaying the hydropathic character of a protein. J. Mol. Biol. 157, 105–132.

Liyanage, V.R.B., Rastegar, M., 2014. Rett syndrome and MeCP2. Neuromolecular Med. 16, 231–264. doi:10.1007/s12017-014-8295-9

Lorenz, R., Bernhart, S.H., Höner Zu Siederdissen, C., Tafer, H., Flamm, C., Stadler, P.F., Hofacker, I.L., 2011. ViennaRNA Package 2.0. Algorithms Mol Biol 6, 26. doi:10.1186/1748-7188-6-26

Lugowski, A., Nicholson, B., Rissland, O.S., 2018. DRUID: a pipeline for transcriptome-wide measurements of mRNA stability. RNA 24, 623–632. doi:10.1261/rna.062877.117

Lugowski, A., Nicholson, B., Rissland, O.S., 2017. Determining mRNA half-lives on a transcriptome-wide scale. Methods 137, 90–98. doi:10.1016/j.ymeth.2017.12.006

Martínez de Paz, A., Ausió, J., 2017. MeCP2, A Modulator of Neuronal Chromatin Organization Involved in Rett Syndrome. Adv. Exp. Med. Biol. 978, 3–21. doi:10.1007/978-3-319-53889-1_1

Mattijssen, S., Arimbasseri, A.G., Iben, J.R., Gaidamakov, S., Lee, J., Hafner, M., Maraia, R.J., 2017. LARP4 mRNA codon-tRNA match contributes to LARP4 activity for ribosomal protein mRNA poly(A) tail length protection. Elife 6, 2173. doi:10.7554/eLife.28889

Melnikov, S., Mailliot, J., Shin, B.-S., Rigger, L., Yusupova, G., Micura, R., Dever, T.E., Yusupov, M., 2016. Crystal Structure of Hypusine-Containing Translation Factor eIF5A Bound to a Rotated Eukaryotic Ribosome. J. Mol. Biol. 428, 3570–3576. doi:10.1016/j.jmb.2016.05.011

Mishima, Y., Tomari, Y., 2016. Codon Usage and 3' UTR Length Determine Maternal mRNA Stability in Zebrafish. Mol. Cell 61, 874–885. doi:10.1016/j.molcel.2016.02.027

Moerke, N.J., Aktas, H., Chen, H., Cantel, S., Reibarkh, M.Y., Fahmy, A., Gross, J.D., Degterev, A., Yuan, J., Chorev, M., Halperin, J.A., Wagner, G., 2007. Small-molecule inhibition of the interaction between the translation initiation factors eIF4E and eIF4G. Cell 128, 257–267. doi:10.1016/j.cell.2006.11.046

Muhlrad, D., Decker, C.J., Parker, R., 1994. Deadenylation of the unstable mRNA encoded by the yeast MFA2 gene leads to decapping followed by 5“-->3” digestion of the transcript. Genes Dev. 8, 855–866.

Nam, J.-W., Rissland, O.S., Koppstein, D., Abreu-Goodger, C., Jan, C.H., Agarwal, V., Yildirim, M.A., Rodriguez, A., Bartel, D.P., 2014. Global Analyses of the Effect of Different Cellular Contexts on MicroRNA Targeting. Mol. Cell 53, 1031–1043. doi:10.1016/j.molcel.2014.02.013

Neymotin, B., Ettore, V., Gresham, D., 2016. Multiple Transcript Properties Related to Translation Affect mRNA Degradation Rates in Saccharomyces cerevisiae. G3 (Bethesda) 6, 3475–3483. doi:10.1534/g3.116.032276

Pechmann, S., Frydman, J., 2013. Evolutionary conservation of codon optimality reveals hidden signatures of cotranslational folding. Nat Struct Mol Bio 20, 237–243. doi:10.1038/nsmb.2466

Presnyak, V., Alhusaini, N., Chen, Y.-H., Martin, S., Morris, N., Kline, N., Olson, S., Weinberg, D., Baker, K.E., Graveley, B.R., Coller, J., 2015. Codon Optimality Is a Major Determinant of mRNA Stability. Cell 160, 1111–1124. doi:10.1016/j.cell.2015.02.029

Radhakrishnan, A., Chen, Y.-H., Martin, S., Alhusaini, N., Green, R., Coller, J., 2016. The DEAD-Box Protein Dhh1p Couples mRNA Decay and Translation by Monitoring Codon Optimality. Cell 167, 122–132.e9. doi:10.1016/j.cell.2016.08.053

Radhakrishnan, A., Green, R., 2016. Connections Underlying Translation and mRNA Stability. J. Mol. Biol. 428, 3558–3564. doi:10.1016/j.jmb.2016.05.025

Rissland, O.S., Norbury, C.J., 2009. Decapping is preceded by 3' uridylation in a novel pathway of bulk mRNA turnover. Nat Struct Mol Bio 16, 616–623. doi:10.1038/nsmb.1601

Rissland, O.S., 2016. The organization and regulation of mRNA-protein complexes. Wiley Interdiscip Rev RNA. doi:10.1002/wrna.1369

Rissland, O.S., Subtelny, A.O., Wang, M., Lugowski, A., Nicholson, B., Laver, J.D., Sidhu, S.S., Smibert, C.A., Lipshitz, H.D., Bartel, D.P., 2017. The influence of microRNAs and poly(A) tail length on endogenous mRNA–protein complexes 1–18. doi:10.1186/s13059-017-1330-z

Sabi, R., Tuller, T., 2014. Modelling the efficiency of codon-tRNA interactions based on codon usage bias. DNA Res. 21, 511–526. doi:10.1093/dnares/dsu017

Saini, P., Eyler, D.E., Green, R., Dever, T.E., 2009. Hypusine-containing protein eIF5A promotes translation elongation. Nature 459, 118–121. doi:10.1038/nature08034

Schmidt, E.K., Clavarino, G., Ceppi, M., Pierre, P., 2009. SUnSET, a nonradioactive method to monitor protein synthesis. Nat Meth 6, 275–277. doi:10.1038/nmeth.1314

Schnall-Levin, M., Rissland, O.S., Johnston, W.K., Perrimon, N., Bartel, D.P., Berger, B., 2011. Unusually effective microRNA targeting within repeat-rich coding regions of mammalian mRNAs. Genome Res. 21, 1395–1403. doi:10.1101/gr.121210.111

Schwanhäusser, B., Busse, D., Li, N., Dittmar, G., Schuchhardt, J., Wolf, J., Chen, W., Selbach, M., 2011. Global quantification of mammalian gene expression control. Nature 473, 337–342. doi:10.1038/nature10098

Shigematsu, M., Honda, S., Loher, P., Telonis, A.G., Rigoutsos, I., Kirino, Y., 2017. YAMAT-seq: an efficient method for high-throughput sequencing of mature transfer RNAs. Nucleic Acids Res. 45, e70. doi:10.1093/nar/gkx005

Shyu, A.B., Greenberg, M.E., Belasco, J.G., 1989. The c-fos transcript is targeted for rapid decay by two distinct mRNA degradation pathways. Genes Dev. 3, 60–72.

Wang, Y., Liu, C.L., Storey, J.D., Tibshirani, R.J., Herschlag, D., Brown, P.O., 2002. Precision and functional specificity in mRNA decay. Proc. Natl. Acad. Sci. U.S.A. 99, 5860–5865. doi:10.1073/pnas.092538799

Webster, M.W., Chen, Y.-H., Stowell, J.A.W., Alhusaini, N., Sweet, T., Graveley, B.R., Coller, J., Passmore, L.A., 2018. mRNA Deadenylation Is Coupled to Translation Rates by the Differential Activities of Ccr4-Not Nucleases. Mol. Cell 70, 1089–1100.e8. doi:10.1016/j.molcel.2018.05.033

Wohlgemuth, I., Brenner, S., Beringer, M., Rodnina, M.V., 2008. Modulation of the rate of peptidyl transfer on the ribosome by the nature of substrates. J. Biol. Chem. 283, 32229–32235. doi:10.1074/jbc.M805316200

Wolf, J., Passmore, L.A., 2014. mRNA deadenylation by Pan2-Pan3. Biochem. Soc. Trans. 42, 184–187. doi:10.1042/BST20130211

Xu, N., Loflin, P., Chen, C.Y., Shyu, A.B., 1998. A broader role for AU-rich element-mediated mRNA turnover revealed by a new transcriptional pulse strategy. Nucleic Acids Res. 26, 558–565.

Yamashita, A., Chang, T.-C., Yamashita, Y., Zhu, W., Zhong, Z., Chen, C.-Y.A., Shyu, A.-B., 2005. Concerted action of poly(A) nucleases and decapping enzyme in mammalian mRNA turnover. Nat Struct Mol Bio 12, 1054–1063. doi:10.1038/nsmb1016

Yang, X., Boehm, J.S., Yang, X., Salehi-Ashtiani, K., Hao, T., Shen, Y., Lubonja, R., Thomas, S.R., Alkan, O., Bhimdi, T., Green, T.M., Johannessen, C.M., Silver, S.J., Nguyen, C., Murray, R.R., Hieronymus, H., Balcha, D., Fan, C., Lin, C., Ghamsari, L., Vidal, M., Hahn, W.C., Hill, D.E., Root, D.E., 2011. A public genome-scale lentiviral expression library of human ORFs. Nat Meth 8, 659–661. doi:10.1038/nmeth.1638

Yu, C.-H., Dang, Y., Zhou, Z., Wu, C., Zhao, F., Sachs, M.S., Liu, Y., 2015. Codon Usage Influences the Local Rate of Translation Elongation to Regulate Co-translational Protein Folding. Mol. Cell 59, 744–754. doi:10.1016/j.molcel.2015.07.018

Zekri, L., Kuzuoğlu-Öztürk, D., Izaurralde, E., 2013. GW182 proteins cause PABP dissociation from silenced miRNA targets in the absence of deadenylation. EMBO J. 32, 1052–1065. doi:10.1038/emboj.2013.44

Zheng, G., Qin, Y., Clark, W.C., Dai, Q., Yi, C., He, C., Lambowitz, A.M., Pan, T., 2015. Efficient and quantitative high-throughput tRNA sequencing. Nat Meth 12, 835–837. doi:10.1038/nmeth.3478

Zufferey, R., Donello, J.E., Trono, D., Hope, T.J., 1999. Woodchuck hepatitis virus posttranscriptional regulatory element enhances expression of transgenes delivered by retroviral vectors. J. Virol. 73, 2886–2892.

